# Targeting *Leishmania infantum* Mannosyl-oligosaccharide glucosidase with natural products: pH-dependent inhibition explored through computer-aided drug design

**DOI:** 10.1101/2024.03.14.585122

**Authors:** Luis Daniel Goyzueta-Mamani, Haruna Luz Barazorda-Ccahuana, Mayron Antonio Candia-Puma, Alexsandro Sobreira Galdino, Ricardo Andrez Machado-de-Avila, Rodolfo Cordeiro Giunchetti, José L. Medina-Franco, Mónica Florin-Christensen, Eduardo Antonio Ferraz Coelho, Miguel Angel Chávez-Fumagalli

## Abstract

Visceral Leishmaniasis (VL) is a serious public health issue, documented in more than ninety countries, where an estimated 500,000 new cases emerge each year. Regardless of novel methodologies, advancements, and experimental interventions, therapeutic limitations, and drug resistance are still challenging. For this reason, based on previous research, we screened natural products (NP) from Nuclei of Bioassays, Ecophysiology, and Biosynthesis of Natural Products Database (NuBBEDB), Mexican Compound Database of Natural Products (BIOFACQUIM), and Peruvian Natural Products Database (PeruNPDB) databases, in addition to structural analogs of Miglitol and Acarbose, which have been suggested as treatments for VL and have shown encouraging action against parasite’s N-glycan biosynthesis. Using computer-aided drug design (CADD) approaches, the inhibitory effect of these NP candidates was evaluated by inhibiting the Mannosyl-oligosaccharide Glucosidase Protein (MOGS) from *Leishmania infantum*, an enzyme essential for the protein glycosylation process, at various pH to mimic the parasite’s changing environment. Also, computational analysis was used to evaluate the Absorption, Distribution, Metabolism, Excretion, and Toxicity (ADMET) profile, while molecular dynamic simulations were used to gather information on the interactions between these ligands and the protein target. Our findings indicated that Ocotillone and Subsessiline have potential antileishmanial effects at pH 5 and 7, respectively, due to their high binding affinity to MOGS and interactions in the active center. Furthermore, these compounds were non-toxic and had the potential to be administered orally. This research indicates the promising anti-leishmanial activity of Ocotillone and Subsessiline, suggesting further validation through *in vitro* and *in vivo* experiments.

## Introduction

Leishmaniasis is a vector-borne disease caused by *Leishmania spp.* protozoan parasites. It is transmitted to humans through the bites of infected female sandflies of *Phlebotomus spp.* in the Old World and *Lutzomyia spp.* in the New World, with distinct clinical symptoms. Leishmaniasis is prevalent in tropical and subtropical regions, with an impact observed in over 90 countries worldwide, particularly in economically disadvantaged areas where sandfly vectors proliferate (Alvar et al., 2012; Akhoundi et al., 2016). This neglected tropical disease manifests in main clinical forms: cutaneous (CL), mucosal (ML), and visceral leishmaniasis (VL); this last one is caused mainly by *L. infantum*, which induces severe symptoms, such as anemia, fever, weight loss, and an enlargement of the liver and spleen, among other internal organs and bone marrow (Ghazanfar and Malik, 2016).

Despite its widespread prevalence, therapeutics for leishmaniasis are limited, with antimonials, single and liposomal amphotericin B, and miltefosine being the most common treatments (Bernal and Coy-Barrera, 2014). However, these drugs often exhibit severe side effects, such as the risk of harming healthy cells and high treatment costs, raising concerns about medication resistance in *Leishmania* parasites. Antimonials’ resistance has compromised the efficacy of these drugs, leading to a pressing need for alternative treatments (Freitas-Junior et al., 2012; Ponte-Sucre et al., 2017). This context highlights the urgency of exploring new avenues for effective and safer treatments for leishmaniasis. On the other hand, developing a vaccine for leishmaniasis has proven to be challenging due to the parasite’s diversity and the host’s immunological response. Despite this challenge, ongoing research in various areas, such as nanomedicine, biomarkers research, and drug repurposing, reveals information on potential treatment alternatives (Kaye et al., 2021).

Researchers are actively working to repurpose current medications and identify accurate biomarkers to monitor treatment response and predict recurrence (Rostami and Khamesipour, 2021; Volpedo et al., 2021). The oral drugs, Miglitol and Acarbose, initially designed for treating type-2 diabetes, are examples of such repurposed drugs. Classified as alpha-glucosidase inhibitors, these drugs function by inhibiting the intestinal digestion and absorption of carbohydrates (Ueno et al., 2015). Notably, Acarbose and Miglitol have demonstrated effectiveness against macrophages infected with *Leishmania*, impacting a key protein in the metabolic network of the N-glycan biosynthesis pathway crucial for the parasite’s survival (Chavez-Fumagalli et al., 2019). Likewise, Acarbose altered the mitochondrial function of *L. infantum*, leading to increased reactive oxygen species (ROS) production and lipid accumulation. This alteration induced a specific production of Th1-type cytokines, reducing parasitism in diverse targeted organs (Costa et al., 2021).

Alternatives for drug development must be researched further to determine their effectiveness. Two critical factors must guide this research: differentiation from the mammalian host and the essential factor of the chosen target for the pathogen’s survival. A parasite’s survival depends on critical metabolic pathways governed by complex chemical mechanisms. Specific enzymes efficiently catalyze a variety of responses in these pathways (Kaye and Scott, 2011). These factors should be carefully considered to identify suitable targets that can effectively disrupt the *Leishmania* survival mechanisms. Research in the field of leishmaniasis focused on uncovering novel biochemical targets associated with various defense mechanisms, including RNA and DNA metabolism, glucose metabolism, sterols, fatty acids, the purine pathway, and nucleotides. The primary objective is identifying targets that new anti-leishmania drugs can effectively damage while ensuring the host’s safety (Jain and Jain, 2018; Raj et al., 2020; Soni and Pratap, 2022).

Consequently, there is an ongoing effort to discover novel inhibitors of proteins and enzymes that function as targets for anti-leishmanial resources. Notably, plant-derived flavonoids, including lupeol, quercetin, and gallic acid, have shown promising activity at comparatively low concentrations against Try-R and Try-S disulfide oxidoreductase enzymes crucial for the detoxification and survival of *Leishmania* (Mehwish et al., 2019). A significant *in vitro* efficacy against *Leishmania* promastigotes and amastigotes and the promising *in vivo* activity of sesquiterpene-related compounds derived from natural sources have been shown (Bernal and Coy-Barrera, 2014). A recent study used molecular docking techniques to analyze numerous terpenoids within the active sites of 24 enzymes from *L. major, L. donovani, L. mexicana,* and *L. infantum* (Ogungbe et al., 2014). Specific enzymatic targets have been studied and tested in this context to develop new drugs to treat leishmaniasis. Examples include Adenine phosphoribosyltransferase (APRT)(Barazorda-Ccahuana et al., 2024), where three furoquinolone alkaloids from the plant *Almeidea rubra* showed APRT inhibitory activity (Ambrozin et al., 2005); N-myristoyltransferase (NMT), in which phenolic compounds, including aurones, chalcones, lignans, and isoflavonoids, showed specific effect and docking activity against NMT (Ogungbe et al., 2014); Dihydroorotate dehydrogenase, where chalcones, flavonoids, monoterpenoids, limonoids, demonstrated anti-leishmanial effect (Ogungbe and Setzer, 2013); and Arginase, in which quercetin epigallocatechin-3-gallate and gallic acid exhibited the most inhibitory effects (dos Reis et al., 2013; Manjolin et al., 2013). Rich natural products (NPs) sources can be found in countries with high biodiversity, such as Peru, which has 1400 described medicinal plant species. These plant-derived NP are cataloged in the PeruNPDB database, where computer-aided drug design (CADD) can be used to discover, develop, and analyze potential drugs and biomolecules against disease (Barazorda-Ccahuana et al., 2023b).

Despite advances in natural and synthetic product research, the precise mechanisms underlying their anti-*Leishmania* mode of action remain unknown. However, environmental factors, such as pH, temperature, and osmotic pressure, may influence the parasite’s life cycle and gene expression. The ideal temperature for parasite species to proliferate as amastigote forms in mammalian hosts is around 37 degrees Celsius (Clos et al., 2022). Similarly, differentiation from promastigotes to amastigotes occurs during macrophage phagocytosis in an acidic environment, which can also occur between 4.5 and 6, although some species can adapt to values between 7 and 7.5. Fluctuations in pH values impact DNA, protein synthesis, and glucose metabolism (Antoine et al., 1990; Garlapati et al., 1999).

In this work, we aim to screen and evaluate compounds contained in three Latin American NP databases, the Nuclei of Bioassays, Ecophysiology, and Biosynthesis of Natural Products Database (NuBBEDB), BIOFACQUIM, and PeruNPDB at different pH levels to identify compounds whose properties and actions are comparable to those of Miglitol and Acarbose. After docking these compounds with the Mannosyl-Oligosaccharide Glucosidase protein (MOGS) enzyme of *L. infantum*, their potential toxic effects were explored with molecular dynamic simulation analyses. This study promotes the research of innovative enzyme inhibitors, consequently expanding the field of medication discovery targeting leishmaniasis, and offers important insights at the molecular level. The insights provided by these analyses suggest the potential for advancing medicines constructed from natural products.

## Methods

### Data collection and structural analogs search

The simplified molecular-input line-entry system (SMILES) (Weininger, 1988) of NPs previously described were retrieved from NuBBEDB online web server (version 2017) (https://nubbe.iq.unesp.br/portal/nubbe-search.html, accessed on 15 August 2023), which contains the information of more than 2,223 NPs and derivatives from Brazilian biodiversity (Pilon et al., 2017); BIOFACQUIM database (https://figshare.com/articles/dataset/BIOFACQUIM_V2_sdf/11312702., accessed on 15 August 2023, which contains the information of 553 NPs from Mexican biodiversity (Pilón-Jiménez et al., 2019); and from the Peruvian Natural Products Database (PeruNPDB) online web server (https://perunpdb.com.pe/, accessed on 15 August 2023), which contains the information of 280 NPs from Peruvian biodiversity (Barazorda-Ccahuana et al., 2023b).

Also, the SMILEs from Acarbose (PubChem CID: 41774) and Miglitol (PubChem CID: 441314) were used for high throughput screening to investigate structural analogs by the SwissSimilarity server (http://www.swisssimilarity.ch/index.php, accessed on 15 September 2023) (Bragina et al., 2022); as the most similar from the commercial types of compounds and the zinc-drug-like compound, library databases were selected, while the combined screening method was chosen for the high throughput screening to achieve the best structural analogs. Default parameters chose threshold values for positivity.

### Molecular properties calculation

The Osiris DataWarrior v05.02.01 software (Sander et al., 2015) was employed to generate the dataset’s structure data files (SDFs). This followed the uploading to the Konstanz Information Miner (KNIME) Analytics Platform (Fillbrunn et al., 2017), where the “Lipinski’s Rule-of-Five” node was employed to calculate physicochemical properties of therapeutic interest—namely: molecular weight (MW), octanol/water partition coefficient (clogP), number of H-bond donor atoms (HBD) and number of H-bond acceptor atoms (HBA)—of the dataset. To generate a visual representation of the chemical space of the dataset for the auto-scaled properties of pharmaceutical interest, the principal component analysis (PCA), which reduces data dimensions by geometrically projecting them onto lower dimensions called principal components (PCs), calculated by the “PCA” KNIME node. Three-dimensional scatter-plot representations were generated for PCA with the Plotly KNIME node.

### Virtual screening

The FASTA sequence of the *L. infantum* Mannosyl-Oligosaccharide Glucosidase protein (MOGS) (ID: A4I3U9) was retrieved from the UniProt database (http://www.uniprot.org/), accessed on 03 October 2023), and subjected to automated modeling in SWISS-MODEL (Biasini et al., 2014). The NP datasets were imported into OpenBabel using the Python Prescription Virtual Screening Tool (Dallakyan and Olson, 2015) and were subjected to energy minimization. PyRx performs structure-based virtual screening by applying docking simulations using the AutoDock Vina tool (Trott and Olson, 2010). The MOGS model was uploaded as a macromolecule, and a thorough search was carried out by enabling the “Run AutoGrid” option, which creates configuration files for the grid parameter’s lowest energy pose, and then the “Run AutoDock” option, which uses the Lamarckian GA docking algorithm. The entire set of modeled 3D models was used as the search space for the study. The docking simulation was then run with an exhaustiveness setting of 20 and instructed only to produce the lowest energy pose. The statistical analysis was done within the GraphPad Prism software version 10.0.2 (232) for Windows from GraphPad Software, San Diego, California, USA, at http://www.graphpad.com. Violin plots were generated for visualization, and the One-way ANOVA followed by Dunnett correction for multiple comparisons test was employed to evaluate the differences between the datasets. The results were considered statistically significant when p<0.05. For the selected compounds, the Tanimoto similarity score was calculated for clustering. The atom-pair-based fingerprints of the compounds were obtained using the “ChemmineR” package (Cao et al., 2008) in the R programming environment (version 4.0.3) (Dalgaard, 2010), and heatmaps were generated for visualization.

### ADMET predictions

The pkCSMonline server, which uses graph-based signatures, was employed for the prediction of the compounds’ ADMET (absorption, distribution, metabolism, elimination, and toxicity) properties (https://biosig.lab.uq.edu.au/pkcsm/, accessed on 14 December 2023)(Pires et al., 2015). The Z-score was calculated for numerical results, while categorical data was converted into binary “Yes” or “No” data. Heatmaps were generated for visualization within the GraphPad Prism software version 10.0.2 (232) for Windows from GraphPad Software, San Diego, California, USA, at http://www.graphpad.com.

### Protonation/deprotonation states by SGCMC

The protein structure input files were established using homology modeling via the Swiss-Model server (https://swissmodel.expasy.org, accessed in July 2023). The original protein was prepared to assess the impact of protonation/deprotonation states of titratable residues at pH 5 and pH 7 using the program developed by Barazorda et al. 2021 (Barazorda-Ccahuana et al., 2021) (accessible online at https://github.com/smadurga/Protein-Protonation). The pKa value for each titratable residue (Asp, Glu, Arg, Lys, and His), along with the C-terminal and N-terminal ends, was determined using Propka v.3 (Rostkowski et al., 2011).

### Molecular docking

The selected compounds underwent design and editing using the Avogadro(x) software, followed by the automatic input generation CHARMM-GUI server (https://www.charmm-gui.org, accessed in September 2023). Before conducting docking, multiple druggable regions of the target protein were delineated, with the optimal druggable site identified using the PASSer program, which locates potential binding pockets within a protein’s structure based on their size, shape, and chemical properties and analyzes these pockets for “druggability”, comparing them to known drug-binding sites for similarity and ranks, up to 100%, the pockets based on their overall potential (https://passer.smu.edu, accessed in September 2023). Molecular docking was conducted using the free web server Dockthor, which performs protein-ligand docking using a genetic algorithm. It prepares the protein and ligand, then iteratively explores different binding poses, scoring them with the MMFF94s force field. The final output is a ranked list of potential binding modes for analysis, docking grids centered on the druggable region. (https://dockthor.lncc.br/v2/, accessed in October 2023).

### Molecular dynamics

The all-atom simulations were conducted using the Gromacs 2023 (Van Der Spoel et al., 2005) (Software package employing the Charmm27 (MacKerell Jr et al., 2000) force field. Each system was enclosed within a cubic box measuring 10 nm³. Explicit water molecules were modeled using the TIP3P (Mark and Nilsson, 2001) water model, and ions were incorporated to achieve system neutrality. The molecular dynamics (MD) simulations comprised three primary stages. Firstly, energy minimization was performed by employing the steep integrator over 200,000 steps. Subsequently, the equilibrium simulation proceeded, wherein restrictions were imposed on the protein-ligand complex for 1 ns within the NVT (number of molecules, volume, and temperature constant) ensemble, followed by an additional restriction in the isobaric-isothermal NPT (number of molecules, pressure, and temperature constant) ensemble for 1 ns. Lastly, the production simulation phase commenced, where restrictions were lifted from the entire system for 100 ns under the isobaric-isothermal NPT ensemble. Estimating binding free energies between the proteins and the ligands was conducted using the Molecular Mechanics/Poisson-Boltzmann Surface Area (MM/PB(GB)SA) methodology. This approach was implemented via the tool called gmx_MMPBSA (Valdés-Tresanco et al., 2021), which was designed to bridge the gap and integrate the extensive functionalities of AmberTools’ MMPBSA.py and associated programs for GROMACS users.

## Results

### Data collection, chemical space visualization, and virtual screening

First, the SMILES from NPs were extracted from three databases: PeruNPDB, BIOFACQUIM, and NuBBEDB. These yielded 280, 553, and 2,223 molecules in total. Utilizing the SwissSimilarity site, 400 and 31 molecules of Miglitol and Acarbose’s structural analogs were found, respectively. A total of 3,487 unique molecules were found, and the duplicate compounds were removed. Figure 1A depicts a visual representation of the chemical space of the data set with 3,487 molecules. The PCA analysis simplifies complex chemical features into three key dimensions. The distribution of the molecules in the plot can indicate the chemical diversity of the generated dataset. A widely scattered plot suggests a more chemically diverse dataset, whereas a plot where most compounds cluster together suggests a less diverse set and might share similar chemical features or potentially similar biological activity.

**Figure 1.**
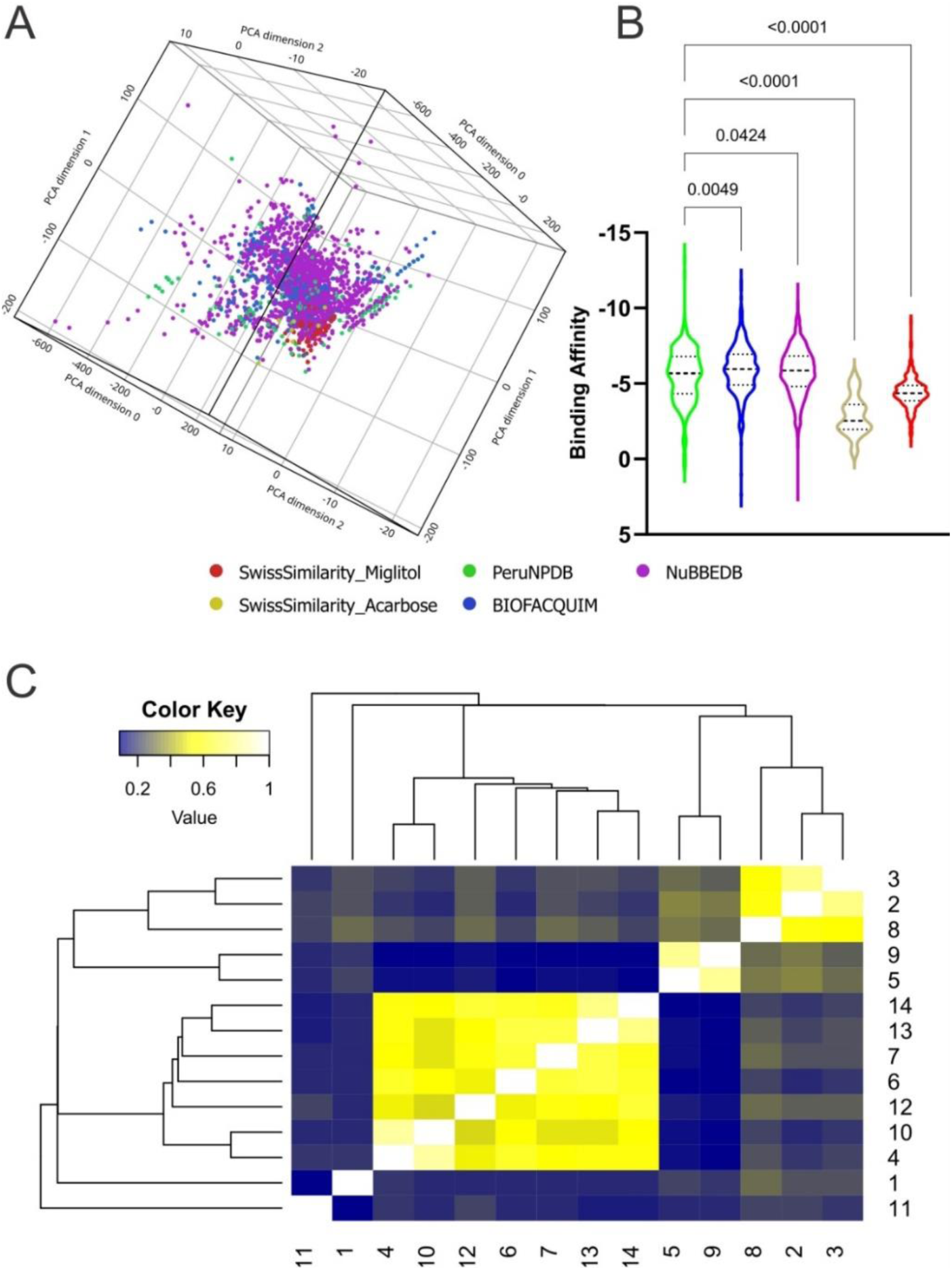
Molecular properties analysis and virtual screening. The chemical space of the generated dataset is represented visually by 3D-PCA **(A)**. binding affinities of NPs against Mannosyl-Oligosaccharide Glucosidase protein **(B)** from the dataset; and heatmap generated with Tanimoto scoring matrix of similar structures among compounds **(C)**.

Subsequently, a virtual screening analysis was conducted on the 3,487 molecules dataset against MOGS (Figure 1B). Among these, eighteen are effectively bound to the protein (Binding affinity ≥10.00 kcal/mol), thus appropriate for additional research. Hence, Tanimoto clustering was used as an efficient tool to screen this set for structural similarity by employing molecular fingerprints that encode relevant structural features and quantify their similarity pairwise. Hierarchical clustering, using the Tanimoto distance (0-1), reveals patterns in the dataset. Clusters containing molecules with high Tanimoto similarity highlight candidates with potentially shared bioactivity (Figure 1C), and observing the ADMET dataset analysis with favorable pharmacokinetic properties, indicating potential bioavailability and reduced systemic toxicity (Figure 2A), three NPs were selected: Subsessiline (PubChem CID: 182033) extracted from *Abuta rufescens* (Swaffar et al., 2012), Ocotillone (PubChem CID: 12313665) extracted from *Cabralea canjerana* (Sarria et al., 2011), and Humilinolide G (PubChem CID: 163100483) extracted from *Swietenia humilis* (Ovalle-Magallanes et al., 2015). Also, the analysis confirmed adherence to Lipinski’s rule of five (molecular weight < 500, logP < 5, more than 5 hydrogen donors and more than 10 hydrogen bond acceptors) increases the likelihood of good oral bioavailability and membrane permeability, absence of PAINS which often act as nonspecific binders in assays, leading to false positives (Dahlin et al., 2015), Brenk alerts avoiding structural motifs flagged helping to minimize potential metabolic liabilities and chemical reactivity issues (Brenk et al., 2008); and Lead likeness characteristics promoting smaller and less complex molecule designs (Teague et al., 1999), suggesting favorable drug-like properties (Figure 2B).

**Figure 2.**
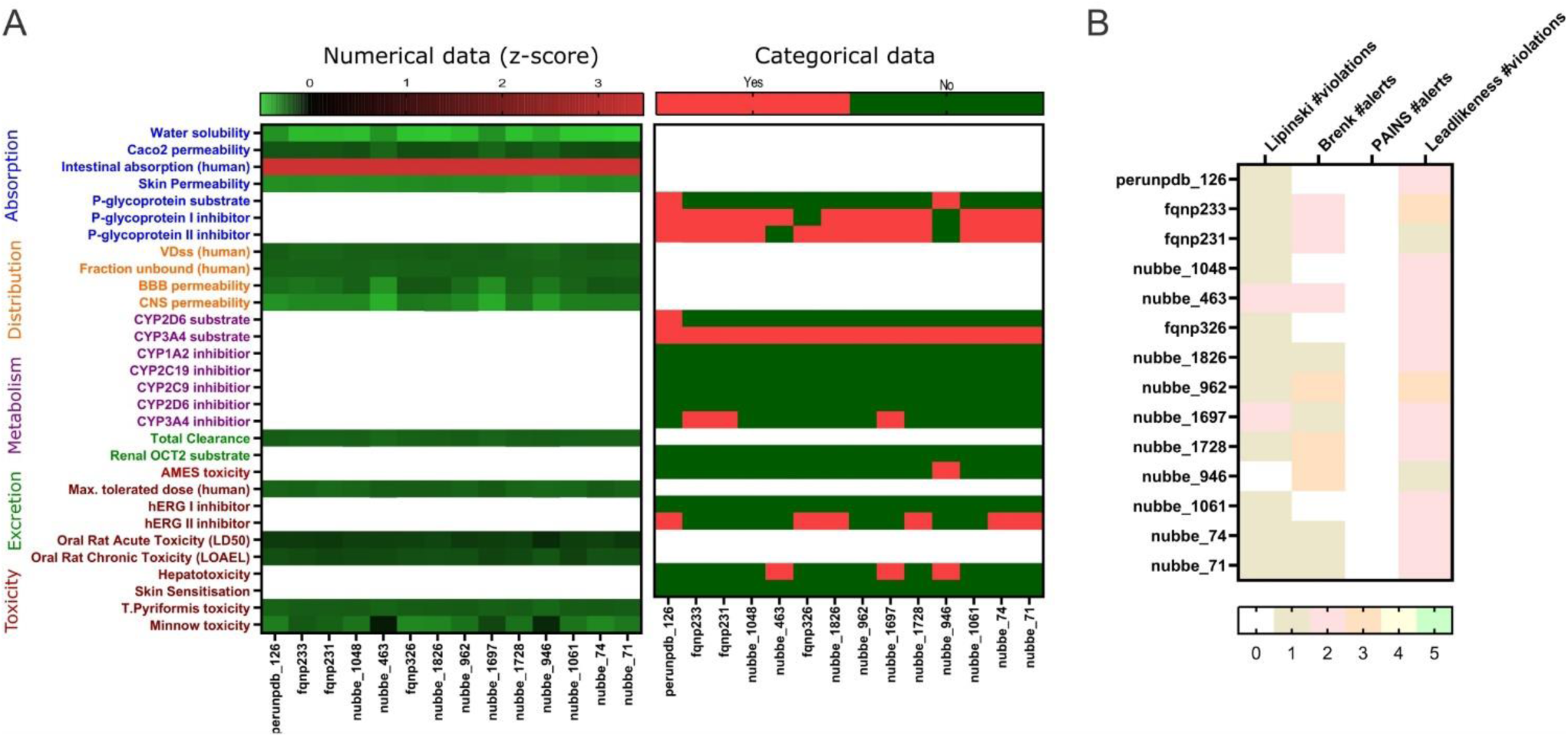
Absorption, Distribution, Metabolism, Excretion (ADMET) prediction heat map of natural products **(A)**; and Lipinski rule of five and PAINS, Brenk alerts, Lead likeness **(B).**

### Molecular dynamics simulation of MOGS

The modeled MOGS protein structure was analyzed using the PASSer software (Figure 3A), which identified 64 druggable regions. The top three druggable pockets were ranked as number 60, 63, and 64, with a druggable probability of 58%, 55%, and 54%, respectively (Figure 3B). The target chosen was pocket number 60 due to the following properties: The druggability score of 0.666, indicates a moderate potential for binding drug-like molecules. Scores closer to 1 suggest higher druggability. The large volume of 1667.223, suggests the pocket can accommodate reasonably sized drug molecules. The high hydrophobicity score of 27.125 indicates a strong preference for non-polar molecules or sections of molecules. The moderate proportion of polar atoms (39.796%) suggests some potential for interaction with polar parts of drug molecules (like hydrogen bond donors or acceptors), and the flexibility score of 0.844 indicates some adaptability. This could allow the pocket to adjust its conformation to fit a drug molecule slightly. The total number of pockets and properties is shown in Table S1.

**Figure 3.**
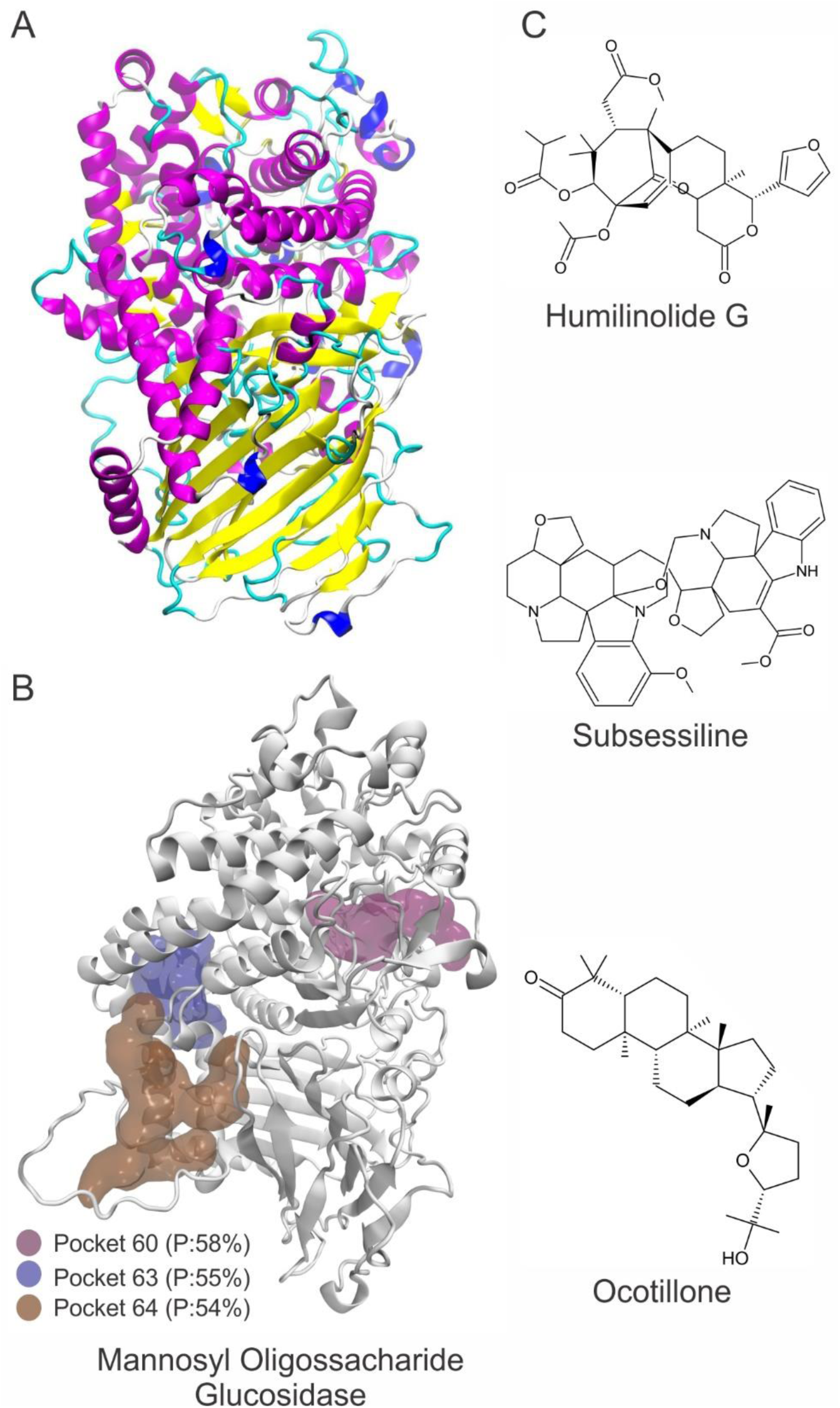
3D Modeled structure of Manossyl Oligosaccharide Glucosidase (MOGS) (left) **(A)** with highlighted most druggable sites (left) **(B)**. The top three natural products found after virtual screening (right) **(C)**: Ocotillone (NuBBEDB), Huminolide G (BIOFACQUIM), and Subsessiline (PeruNPDB). P=Druggable probability.

The thermodynamic parameters were calculated and computed with a 100 ns NPT simulation snapshot. Analysis of the backbone involved assessing the root mean squared deviation (RMSD) and root mean squared fluctuations (RMSF) per residue. Furthermore, the radius of gyration (RG) was assessed, quantifying the mass distribution dispersion concerning the central axis. The compactness of a protein, crucial for its folding rate, was directly linked to RG, especially when computed using advanced computational methods (Lobanov et al., 2008). To ascertain the accessible surface area of solvent molecules on the protein, the solvent-accessible surface area (SASA) was analyzed. Conformational changes occur in proteins in response to external stresses, such as the binding of a foreign substance like a medication; these changes increase the solubility of hydrophobic residues.

In general, the average RG and SASA values of the study indicated that proteins linked to pharmaceutical substances had small conformational changes, most likely as a result of ligands occupy ing their active sites (Durham et al., 2009). The tetrameric conformation of the modeled MOGS exhibited a stable behavior throughout the 100 ns snapshot of MDS conducted at pH 7. It showed a more stable behavior than 5, as shown in Figure 4. The RMSD values of the systems performed to verify the similarity between a protein-bound and not-bound ligand for MOGS receptor were as follows: MOGS (pH 5: 0.23 nm and pH 7: 0.22 nm), MOGS:Humilinolide G (pH 5: 0.21 nm and pH 7: 0.31 nm), MOGS:Ocotillone acid (pH 5: 0.28 nm and pH 7: 0.26 nm), and MOGS:Subsessiline (pH 5: 0.25 nm and pH 7: 0.27 nm). On the other hand, the MOGS: Ocotillone system exhibits low fluctuations in loop peaks as measured by the RMSF per residue, with low peaks at pH 5 at 0.1065, 0.1213, and 0.5732 nm. Meanwhile, the MOGS: Subsessiline system shows low peaks at pH 7 at 0.0528, 0.0564, 0.0689, 0.0694, 0.0734, 0.1197, 0.1380, and 0.2151 nm. The greater RG values obtained from Ocotillone at pH 5 (2.90 nm) and Subsessiline at pH 7 (2.89 nm) showed that the MOGS structure was less compact. In the same way, the average SASA value of Ocotillone at pH 5 (381.80 nm^2^) and Subsessiline at pH 7 (378.86 nm^2^) was higher in comparison to the remaining compounds, indicating lower compactation of MOGS.

**Figure 4.**
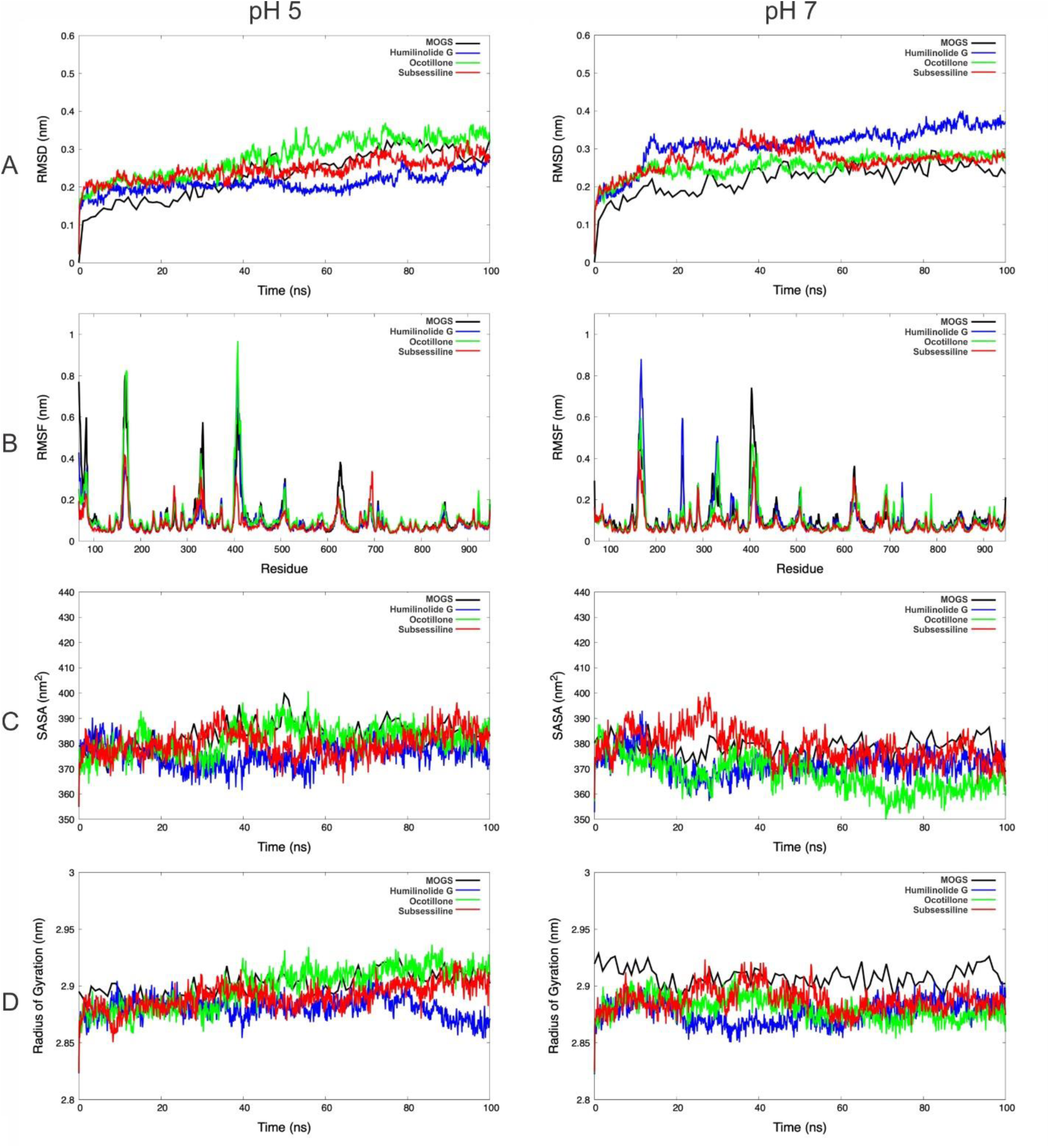
Representation of DM trajectories over 100 ns for Mannosyl-Oligosaccharide Glucosidase, Humilinolide G, Ocotillone, and Subsessiline. The figures on the left and right of the panel above were performed at pH 5 and pH 7, respectively. **(A)** Root mean squared deviation (RMSD); **(B)** root mean squared fluctuations (RMSF); **(C)** solvent-accessible surface area (SASA); and **(D)** radius of gyration (RG).

### Analysis of Protein-Ligand Binding Affinities with Molecular Mechanics Generalized Born Surface Area Calculations (MM/GBSA)

The estimation of the binding free energy (ΔG) is made possible by the utilization of continuum solvation implicit models in conjunction with Molecular Mechanics Poisson–Boltzmann Surface Area (MM/PBSA) and Molecular Mechanics Generalized Born Surface Area (MM/GBSA) methods. Through the examination of multiple conformations retrieved from the last 100 frames of the MD simulations, the ΔG values were determined in which the most suitable results were the systems MOGS:Ocotillone at pH 5 (-12.96 Kcal·mol^−1^) and MOGS:Subsessiline at pH 7 (-18.67 Kcal·mol^−1^) suggesting them as the most suitable candidates among the three chosen initially. The estimated phase-gas binding free energy (ΔGgas) also provided the highest energy contribution for this system, with values of -47.62 and -32.98 Kcal·mol^−1^, respectively.

The data shown in Table 2 emphasizes the significant effect of van der Waals energies alongside electrostatic and generalized born energies. The analysis at two different pH levels can positively or negatively influence the protein/ligand binding due to its effect on hydrophobic interactions. In this context, the electrostatic energies (ΔEele) of Subsessiline, among others, contributed more positively to this binding at a pH of 7 than a pH of 5. The aforementioned pattern was similarly noted in the measurement of solvation energies (ΔGsolv), where negative values indicate an unfavorable contribution to the protein-ligand interaction. Based on the 2D interaction summary represented in Figure 5, it is evident that, at pH 7, the ligand Subsseline effectively replicated a critical hydrogen bonding interaction with specific amino acid residue ARG710.

**Figure 5.**
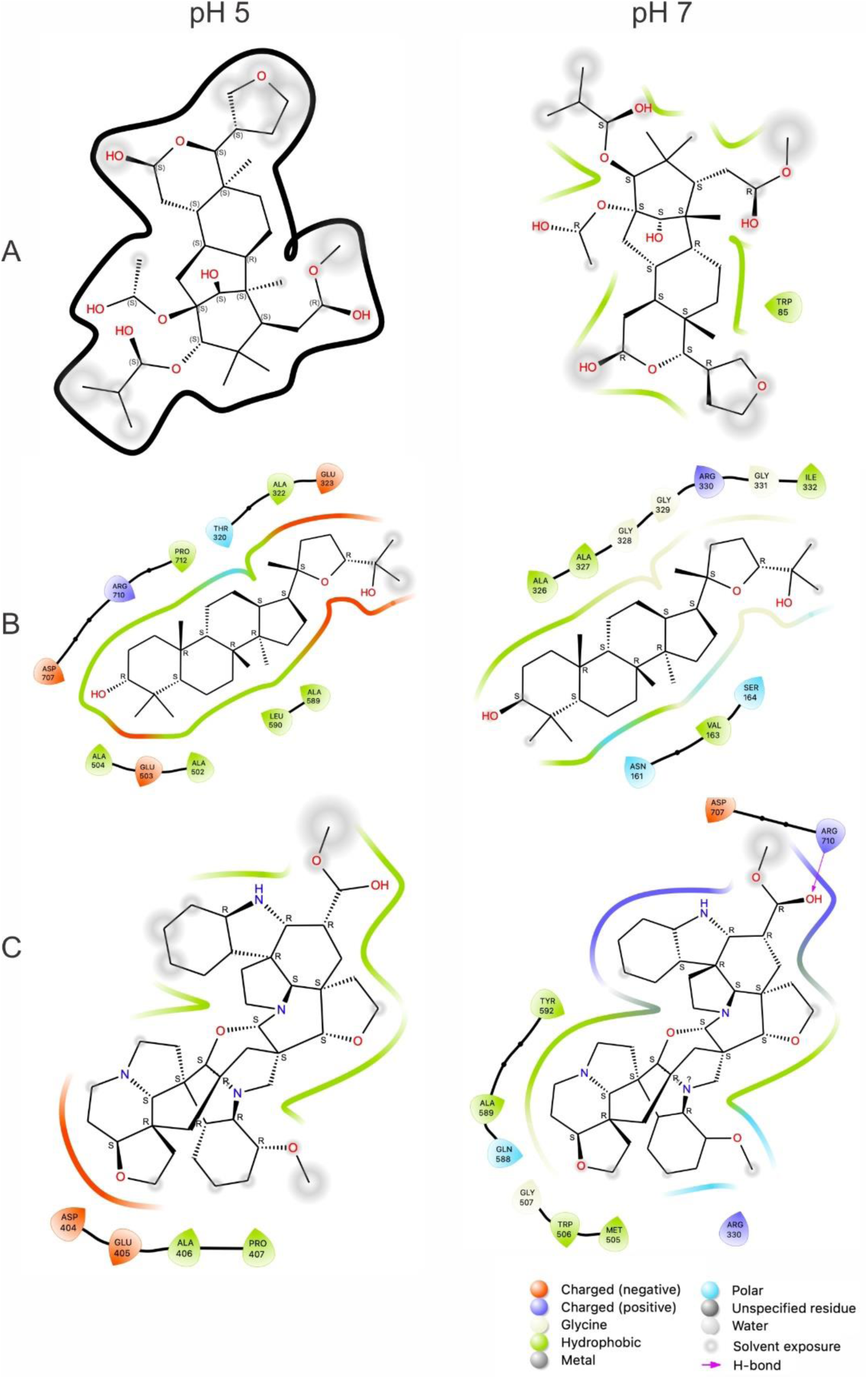
2D representation of the top inhibitors with Mannosyl-Oligosaccharide Glucosidase and compounds computed from 100 ns MD simulation. **(A)** Humilinolide G, **(B)** Ocotillone, and **(C)** Subsessiline. The bottom legends identify the types of interaction and bonds with their corresponding color codes. Thick black line in section A indicates no binding between NP and MOGS.

**Table 1.** RMSD, SASA, and RG, average values of the top three targets: Humilidoline G, Ocotillone and Subsessiline, based on 100 ns of MD simulations.

| Structure | pH 5 |  |  |  |  |  | pH 7 |  |  |  |  |  |
| --- | --- | --- | --- | --- | --- | --- | --- | --- | --- | --- | --- | --- |
|  | RMSD (nm) |  | SASA (nm<sup>2</sup>) |  | RG (nm) |  | RMSD (nm) |  | SASA (nm<sup>2</sup>) |  | RG (nm) |  |
|  | Average | DesV | Average | DesV | Average | DesV | Average | DesV | Average | DesV | Average | DesV |
| MOGS | 0.23 | 0.07 | 382.84 | 5.57 | 2.90 | 0.01 | 0.22 | 0.04 | 378.32 | 4.05 | 2.91 | 0.01 |
| Humilidoline G | 0.21 | 0.02 | 375.63 | 4.96 | 2.88 | 0.01 | 0.31 | 0.05 | 371.76 | 4.95 | 2.88 | 0.01 |
| Ocotillone | 0.28 | 0.05 | 381.80 | 5.70 | 2.90 | 0.02 | 0.26 | 0.03 | 367.60 | 6.64 | 2.88 | 0.01 |
| Subsessiline | 0.25 | 0.03 | 379.83 | 5.32 | 2.89 | 0.01 | 0.27 | 0.03 | 378.86 | 6.39 | 2.89 | 0.01 |

**Table 2.**
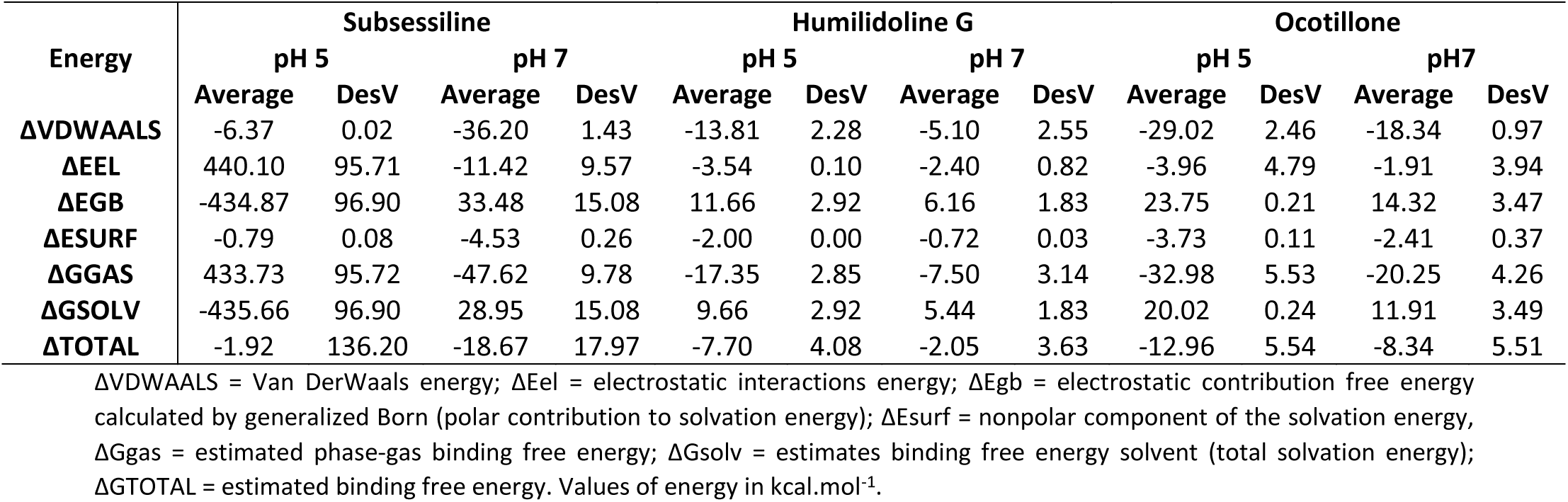
Average value of the binding free energy determined by the MM/PBSA method.

| Energy | Subsessiline |  |  |  | Humilidoline G |  |  |  | Ocotillone |  |  |  |
| --- | --- | --- | --- | --- | --- | --- | --- | --- | --- | --- | --- | --- |
|  | pH 5 |  | pH 7 |  | pH 5 |  | pH 7 |  | pH 5 |  | pH7 |  |
|  | Average | DesV | Average | DesV | Average | DesV | Average | DesV | Average | DesV | Average | DesV |
| **ΔVDWAALS** | -6.37 | 0.02 | -36.20 | 1.43 | -13.81 | 2.28 | -5.10 | 2.55 | -29.02 | 2.46 | -18.34 | 0.97 |
| **ΔEEL** | 440.10 | 95.71 | -11.42 | 9.57 | -3.54 | 0.10 | -2.40 | 0.82 | -3.96 | 4.79 | -1.91 | 3.94 |
| **ΔEGB** | -434.87 | 96.90 | 33.48 | 15.08 | 11.66 | 2.92 | 6.16 | 1.83 | 23.75 | 0.21 | 14.32 | 3.47 |
| **ΔESURF** | -0.79 | 0.08 | -4.53 | 0.26 | -2.00 | 0.00 | -0.72 | 0.03 | -3.73 | 0.11 | -2.41 | 0.37 |
| **ΔGGAS** | 433.73 | 95.72 | -47.62 | 9.78 | -17.35 | 2.85 | -7.50 | 3.14 | -32.98 | 5.53 | -20.25 | 4.26 |
| **ΔGSOLV** | -435.66 | 96.90 | 28.95 | 15.08 | 9.66 | 2.92 | 5.44 | 1.83 | 20.02 | 0.24 | 11.91 | 3.49 |
| **ΔTOTAL** | -1.92 | 136.20 | -18.67 | 17.97 | -7.70 | 4.08 | -2.05 | 3.63 | -12.96 | 5.54 | -8.34 | 5.51 |

ΔVDWAALS = Van DerWaals energy; ΔEel = electrostatic interactions energy; ΔEgb = electrostatic contribution free energy calculated by generalized Born (polar contribution to solvation energy); ΔEsurf = nonpolar component of the solvation energy, ΔGgas = estimated phase-gas binding free energy; ΔGsolv = estimates binding free energy solvent (total solvation energy); ΔGTOTAL = estimated binding free energy. Values of energy in kcal.mol<sup>-1</sup>.

## Discussion

VL disproportionately impacts economically disadvantaged communities in tropical and subtropical areas, with an estimated 500,000 new cases annually (Alvar et al., 2012; Africa and Asia, 2018). Sudan, Ethiopia, Brazil, Kenya, Somalia, and South Sudan represent most cases (Hotez et al., 2020). If acute and untreated, the death rate associated with VL exceeds 95% (Rodrigues Monteiro et al., 2023). Treatment options are hampered by difficulties, such as toxicity, prolonged administration, and the emergence of drug resistance; furthermore, the unavailability of a vaccine hinders prevention efforts. The urgent necessity for improved diagnostics, vector control strategies, novel and safer medications, an effective vaccine, and an emphasis on addressing the social and economic disparities that sustain the transmission of this neglected tropical disease is highlighted by the challenges raised by VL (De Rycker et al., 2018).

NPs, which have enhanced bioactive potential and structural diversity, are a potentially promising source of possible anti-leishmanial compounds. Implementing CADD techniques to repurpose NPs may accelerate drug discovery and improve affordability. CADD makes virtual screening against *Leishmania* targets possible, enhancing potential NP scaffolds (Barazorda-Ccahuana et al., 2023a, 2024; Luna et al., 2023). Examples of natural compounds that have shown anti-leishmanial activity comprise alkaloids (like berberine and piperine) (Sakyi et al., 2021), flavonoids (such as quercetin and rutin) (Mehwish et al., 2021), and terpenes (like limonene and eugenol) (Almeida-Bezerra et al., 2022). Despite the ongoing difficulties associated with compound isolation and validation, integrating NPs research and CADD, including chemoinformatics (Medina-Franco and Saldívar-González, 2020), emerges as a persuasive approach to address the increasing demand for novel and efficacious treatments for VL.

Due to the promising results obtained in our previous research (Chavez-Fumagalli et al., 2019; Costa et al., 2021), we utilized and modeled MOGS (formerly categorized as a hypothetical protein) as a potential target and receptor in this study to evaluate the effect of NPs (Figure 3A). Notably, in early genome sequencing endeavors, numerous computationally predicted open reading frames (ORFs) encoded proteins lacking functional annotation, called hypothetical proteins (Amatore et al., 2020). An ORF that exhibited sequence homology to enzymes in the Glycoside Hydrolase (GH) family attracted attention. Thorough biochemical characterization confirmed its activity as a MOGS (Nairn and Moremen, 2014). MOGS is crucial in the endoplasmic reticulum (ER) N-linked glycosylation pathway (Kuribara and Totani, 2022). The hydrolytic removal of the terminal α-1,2-linked glucose residue from the Glc3Man9GlcNAc2 oligosaccharide precursor is catalyzed at that location. This critical trimming step impacts glycoprotein trafficking, ER quality control mechanisms, and protein folding (Ito et al., 2020; Kuribara and Totani, 2021; Ninagawa et al., 2021).

*Leishmania*, parasites exhibit N-glycosylation patterns distinct from their mammalian hosts. MOGS is involved in these pathways, and research suggests that it contributes to the biosynthesis of Lipophosphoglycan (LPG), an essential defense mechanism for the promastigote stage of *Leishmania* that guarantees the parasite’s survival within the sandfly vector (Lodge and Descoteaux, 2005). Additionally, interactions between macrophage receptors and MOGS-processed parasite surface glycoproteins may contribute to *Leishmania’s* invasion of the host cell (Olafson et al., 1990). Thus, identifying druggable pockets within proteins is a crucial tool in drug design, enabling targeted modulation of protein activity for therapeutic development. Our analysis of the modeled MOGS structure revealed promising druggable pockets, with pocket number 60 demonstrating particularly favorable characteristics (Figure 3B). The studies conducted in this work helped to confirm this pocket’s capability to bind to NPs, highlighting its potential to influence the biological activity of this glucosidase.

Consequently, the current study aimed to use CADD approaches to identify NPs from the NuBBEDB, BIOFACQUIM, and PeruNPDB databases, as well as Miglitol and Acarbose analogs that may have antileishmanial and anti-MOGS properties. At the outset, we distinguished three compounds, namely Ocotillone (PubChem CID 12313665), Subsessiline (PubChem CID 182033), and Humilinolide G (PubChem CID 163100483) (Figure 3C), which stabilized the MOGS structure with an average RMSD higher than that of the protein without compounds. The relevance of certain N-glycosylation processes on the biology of *Leishmania* highlights the promise of MOGS as a prospective target for anti-leishmanial drugs (Naderer et al., 2004). Peptide-specific inhibitors, such as those evaluated here, can potentially interfere with critical N-glycan production pathways. A lack of homology between *Leishmania* MOGS and its human counterpart can significantly enhance the drug’s specificity, thereby reducing toxicity toward the host (Chavez-Fumagalli et al., 2019). Although these NPs have significant potential for drug development, their effectiveness within macrophages and MOGS can be significantly affected by the environment. An effective method for analyzing this effectiveness was using molecular docking and dynamic simulations. These techniques can extensively consider crucial factors such as pH levels, as they possess the potential to drastically modify the charge distribution, conformation, and binding characteristics of the NPs.

It is important to note that *Leishmania* has a unique lifecycle encompassing two stages. The promastigote stage thrives within its sandfly host, facing conditions ranging from slightly acidic to alkaline. Upon entering a mammalian host, it transforms into the amastigote stage, adapted to survive within the acidic environment of the phagolysosomes of macrophages (Rosenzweig et al., 2008). Understanding insight into the processes by which *Leishmania* recognizes, adapts to, and utilizes these significant fluctuations in pH is critical to unraveling its interesting life cycle and discovering prospective treatment targets. Although several organisms would perish in the acidic environment of the phagolysosome of a host cell, *Leishmania* has developed ways that allow it to survive and develop in this situation. An examination of the impacts of pH reveals that the parasite possesses a resistance mechanism to acid levels, responding through stress-response pathways and specific transporters to maintain intracellular balance (McConville and Naderer, 2011). Moreover, *Leishmania* prevents the usual phagosome maturation process, ensuring a safe environment within the host cell and avoiding exposure to the lysosomal damaging enzymes and highest acidity (Podinovskaia and Descoteaux, 2015).

Significantly, *Leishmania* utilizes the acidic environment (pH 4.5 - 5.5) of phagolysosomes as an important trigger for its development. Research findings indicate that the parasite may have pH-sensing systems that initiate its transformation from external promastigotes to intracellular amastigotes (Pal et al., 2017). Significant modifications in gene expression occur during this transition, leading to the synthesis of surface chemicals and proteins optimized for the survival of amastigote cells (Cohen-Freue et al., 2007). Likewise, the function of MOGS, in this pH-dependent transition, is an exciting topic for further studies. Consequently, considering this information, we performed a molecular dynamic simulation at two different pH levels, 5 and 7, to evaluate the effects of this parameter (Figure 4).

At pH 5, also shown by the values in Table 1. significant conformational changes in the protein were found in interaction with Ocotillone, as indicated by an RMSD value of 0.28±0.05 nm, which was higher than the value of the protein in the absence of ligands. Subsessiline subsequently followed with an RMSD of 0.25±0.03 nm. Humidilidoline G, in comparison to the aforementioned, produced the most stable protein structure, as indicated by its average RMSD value of 0.21±0.02. This is because, in this system, the active center does not retain the compound for an extended time at pH 5. Concerning the solvent-accessible solvent area (SASA) and radius of gyration (RG), Ocotillone and Subsessiline have more effective results than Humilidoline G. The stability of the compound related to MOGS at pH 5 was Ocotillone>Subsessiline>Humidilidoline G. Additionally, our research revealed the catalytic dyad for MOGS, a key component of the protein binding site, composed of hydrophobic bonds with Ocotillone ALA 589 (0.1213 nm), ARG 710 (0.1065 nm), and PRO 172 (0.5732 nm) (Figure 6A).

**Figure 6.**
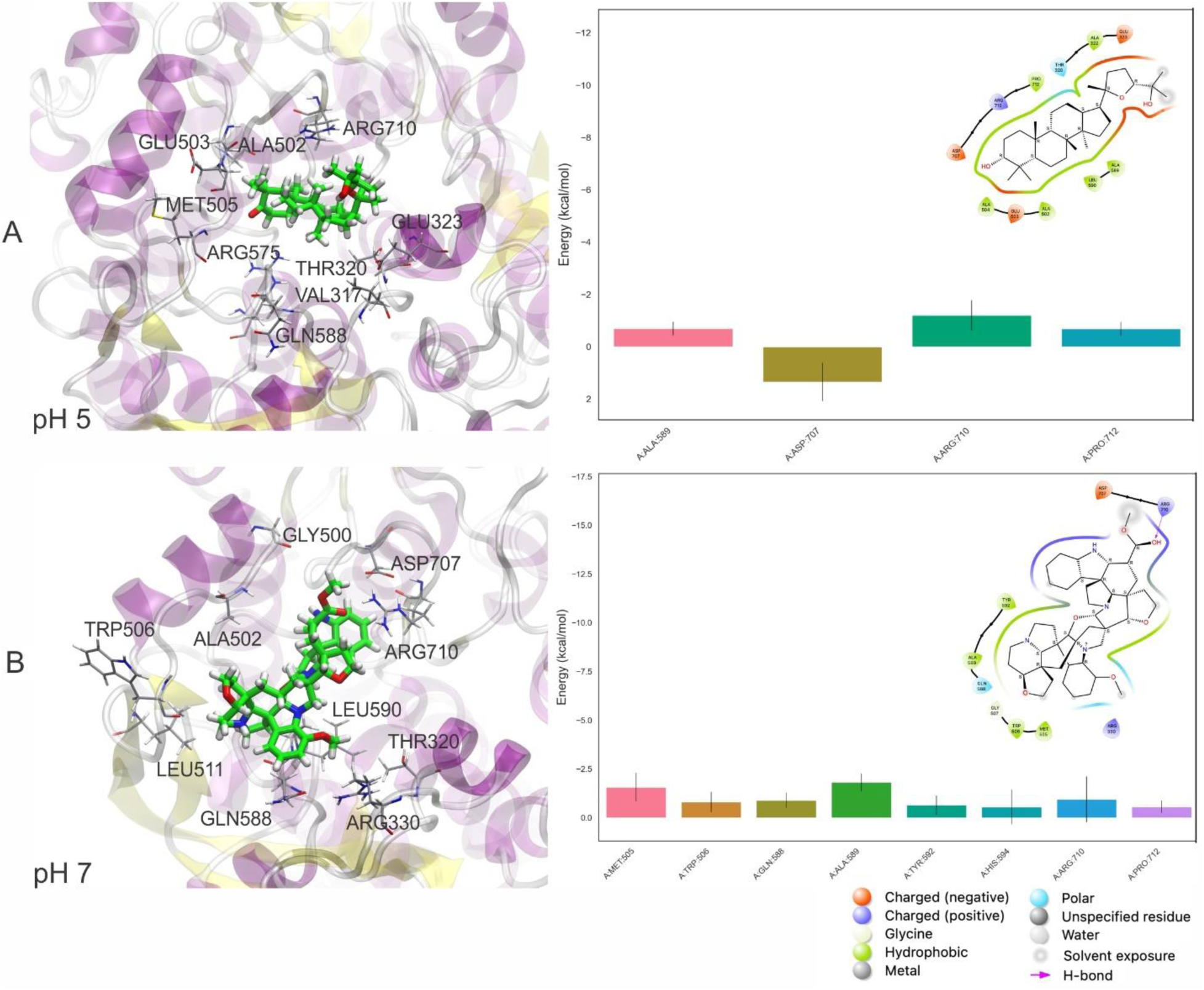
Pictorial 3D (Left) and 2D (right) representation of the best binding free energy between Ocotillone **(A)** and Subsessiline **(B)** at pH 5 and 7, respectively; and Mannosyl-Oligosaccharide Glucosidase (MOGS), computed from the last 10 ns (100 frames) of 100 ns MD simulation.

At pH 7, the compounds’ behavior increases the therapeutic target’s conformational changes. Notably, Humilidoline G and Ocotillone exhibited short retention at the active center within these systems (10 ns of 100 ns). Meanwhile, Subsessiline was the compound that remained in its position in the active site after MD concluded. Nevertheless, the proteins binding to ligands exhibited more significant oscillations in the systems’ RMSD values than those without ligands. In systems involving Humilidoline G and Ocotillone, the solvent has reduced access to the protein surface, as indicated by the SASA value. The stability of the compound related to MOGS at pH 7. was Subsessiline>Humidilidoline G> Ocotillone. The catalytic dyad for MOGS (Figure 6B)., composed of hydrophobic bonds with Subsessiline MET 505 (0.1197 nm), TRP 506 (0.1380 nm), GLN 588 (0.0689 nm) ALA 589 (0.0694 nm) TYR 592 (0.0564 nm) HIS 594 (0.0528 nm) and PRO 172 (0.2151 nm), and hydrogen bond ARG 710 (0.0734 nm) (Figure 5C). The specificity and directionality of hydrogen bonds enhance drug selectivity, reducing off-target effects. These bonds also stabilize the drug-target complex, potentially influencing a drug’s residence time and improving its overall efficacy. Unfortunately, we could not find literature discussing the role of specific amino acid residues in this protein’s conformation, highlighting its novelty and the need for further research.

Ocotillone and Subsessiline might inhibit protein folding in *Leishmania*’s transition from promastigote to amastigote stage, offering a potential treatment for Leishmaniasis. Focusing on this critical phase, these NPs could limit the parasite’s capacity to settle within the macrophage, preventing its survival and disease development. Moreover, these NPs demonstrate stable performance across different pH levels in the macrophage’s phagolysosome, indicating their efficacy throughout the infection process. This stability is crucial because the parasite effectively takes advantage of these pH changes to control the phagolysosome environment in its favor. Furthermore, evaluating ADMET (absorption, distribution, metabolism, excretion, and toxicity) characteristics and adherence to Lipinski’s Rule of Five may predict low toxicity linked to these natural molecules. These assays provide an important initial evaluation of a drug candidate’s drug-likeness and potential for absorption through the digestive system, including parameters such as molecular size, lipophilicity, and hydrogen bonding potential.

Ocotillone, a naturally occurring compound isolated from *Cabralea canjerana*, a native tree in Brazil, has a promising potential for a wide range of biological applications. It shows significant antifeedant efficacy against *Spodoptera litura* in insect bioassays, as evidenced by a Percentage Feeding Index (PFI) that is comparable to that of well-established biopesticides such as Limonin, Azadiradione, and Epoxyazadiradione (Sarria et al., 2011). Furthermore, Ocotillone shows considerable *in vitro* anticancer activity when tested against cancer cell lines MCF-7 (breast), NCI-H460 (lung), and A375-C5 (melanoma). Also, it synergises with colchicine, combating multidrug resistance (MDR) in cancer treatment (M Cazal et al., 2010). Moreover, dichloromethane extracts derived from closely related species (e.g., *P. grandiflorus*) containing Ocotillone in its composition exhibit a significant inhibitory effect against the fungus *Leucoagaricus gongylophorus* (de Souza et al., 2005).

Subsessiline, on the other hand, was initially isolated from the native tree of the Peruvian Amazon, *Abuta rufescens* (Swaffar et al., 2012), and demonstrated *in vitro* efficacy against *Plasmodium falciparum*. Although the precise efficacy of the substance was not disclosed, it exhibited an IC50 ranging from 4 to 40 micrograms per milliliter, inhibiting the *in vitro* proliferation of parasites. Notably, this extract also inhibited the parasite’s capacity to transform hazardous heme into innocuous hemozoin crystals, indicating the possibility of a second mechanism of action (Ruiz et al., 2011). Roumy et al., 2007, also investigated this effect and evaluated the antiplasmodial activity of *Abuta rufescens*, which is historically utilized by the Shipibo-Conibo community in Peru. The activity of the leaves and bark extracts against *P. falciparum* was found to be good to moderate, as indicated by IC50 values ranging from 2.3 to 7.9 g/mL. This discovery is consistent with antiplasmodial alkaloids previously documented in the stem (such as isoquinoline, azafluoranthene, and oxoaporphine), supporting the traditional use of *A. rufescens* as a therapy for malaria.

Examining pH dynamics in *Leishmania* makes a significant contribution that reaches beyond basic comprehension of the parasite’s biology and pathology. It provides valuable insights into potential treatment pathways, making room for novel strategies to overcome a disease affecting millions of people worldwide. Researchers perform phagosome infection models and *in vitro* pH manipulations with great attention to detail to analyze pH sensing, adaptability, and differentiation mechanisms. Critical molecular targets, such as MOGS, for developing novel drugs could be proposed by revealing these systems. Furthermore, it is significant to understand the impact of pH on the absorption, stability, and efficacy of drugs to develop highly effective anti-leishmanial substances. Notably, therapeutic approaches could potentially achieve a synergistic effect by concurrently targeting *Leishmania*’s pH regulating systems and inhibitors of differentiation processes, including those dependent on MOGS-dependent N-glycosylation. To the best of our knowledge, no studies have been published on the pharmacological potential activity of Ocotillone and Subsessiline against Leishmania. While considering the pH variations, the MOGS as a target and comparing and validating this catalytic dyad is still challenging.

## Conclusions

This study aimed to identify *L. infantum* inhibitors from NP scaffolds through *in silico* analysis and database mining. Based on similarity analysis of NPs and existing data in three major public NP databases (NuBBEDB, BIOFACQUIM, and PERUNPDB). Ocotillone (PubChem CID: 12313665) and Subsessiline (PubChem CID: 182033) were evaluated at pH 5 and 7, respectively, and emerged as the most promising candidates. *In silico* docking analyses indicated a favorable affinity for the *L. infantum* MOGS enzyme that could suggest antileishmanial activity. It is worth mentioning that the compounds showed no predicted toxicity and were shown *in silico* to be compatible with oral administration. Additional *in vitro* and *in vivo* experiments are necessary to validate the possible therapeutic potential of Ocotillone and Subsessiline in the treatment against VL, as suggested by the results described here.

## Author Contributions

LDGM, MACP, ASG, RAMA, RCG, JLMF, MFC, and EAFC helped conceptualize and review the study. LDGM and HLBC were the primary authors of the manuscript, with substantial input in the interpretation of findings, critical feedback, and assistance in editing from MACF, MFC, and EAFC. All authors contributed to the creation of the final manuscript.

