## SUPPLEMENTARY MATERIAL for "Targeting *Leishmania infantum* Mannosyl-oligosaccharide glucosidase with natural products: pH-dependent inhibition explored through computer-aided drug design"

**Table S1. PASSer predictions for allosteric site probabilities in MOGS protein pockets.**

| Pocket Number | Parameter | Amount |
|---|---|---|
| Pocket 1 | | |
| | Score | 0.118 |
| | Druggability Score | 0.006 |
| | Number of Alpha Spheres | 50.000 |
| | Total SASA | 130.671 |
| | Polar SASA | 69.082 |
| | Apolar SASA | 61.589 |
| | Volume | 488.359 |
| | Mean local hydrophobic density | 5.250 |
| | Mean alpha sphere radius | 3.932 |
| | Mean alp. sph. solvent access | 0.707 |
| | Apolar alpha sphere proportion | 0.160 |
| | Hydrophobicity score | -3.000 |
| | Volume score | 3.308 |
| | Polarity score | 10.000 |
| | Charge score | -2.000 |
| | Proportion of polar atoms | 51.351 |
| | Alpha sphere density | 6.177 |
| | Cent. of mass - Alpha Sphere max dist | 14.690 |
| | Flexibility | 0.922 |
| Pocket 2 | | |
| | Score | 0.084 |
| | Druggability Score | 0.002 |
| | Number of Alpha Spheres | 20.000 |
| | Total SASA | 18.505 |
| | Polar SASA | 6.429 |
| | Apolar SASA | 12.076 |
| | Volume | 96.898 |
| | Mean local hydrophobic density | 7.000 |
| | Mean alpha sphere radius | 3.624 |
| | Mean alp. sph. solvent access | 0.414 |
| | Apolar alpha sphere proportion | 0.400 |
| | Hydrophobicity score | 9.889 |

| | | |
|---|---|---|
| | Volume score | 4.556 |
| | Polarity score | 5.000 |
| | Charge score | 2.000 |
| | Proportion of polar atoms | 40.000 |
| | Alpha sphere density | 1.723 |
| | Cent. of mass - Alpha Sphere max dist | 3.745 |
| | Flexibility | 0.897 |
| Pocket 3 | | |
| | Score | 0.074 |
| | Druggability Score | 0.001 |
| | Number of Alpha Spheres | 19.000 |
| | Total SASA | 11.554 |
| | Polar SASA | 4.308 |
| | Apolar SASA | 7.246 |
| | Volume | 83.399 |
| | Mean local hydrophobic density | 8.000 |
| | Mean alpha sphere radius | 3.606 |
| | Mean alp. sph. solvent access | 0.454 |
| | Apolar alpha sphere proportion | 0.474 |
| | Hydrophobicity score | -5.000 |
| | Volume score | 4.500 |
| | Polarity score | 6.000 |
| | Charge score | 1.000 |
| | Proportion of polar atoms | 33.333 |
| | Alpha sphere density | 1.325 |
| | Cent. of mass - Alpha Sphere max dist | 2.830 |
| | Flexibility | 0.941 |
| Pocket 4 | | |
| | Score | 0.066 |
| | Druggability Score | 0.541 |
| | Number of Alpha Spheres | 53.000 |
| | Total SASA | 151.351 |
| | Polar SASA | 48.703 |
| | Apolar SASA | 102.648 |
| | Volume | 648.796 |
| | Mean local hydrophobic density | 20.857 |
| | Mean alpha sphere radius | 3.872 |
| | Mean alp. sph. solvent access | 0.530 |
| | Apolar alpha sphere proportion | 0.528 |
| | Hydrophobicity score | -4.857 |
| | Volume score | 4.857 |
| | Polarity score | 10.000 |

| | | |
|---|---|---|
| | Charge score | 7.000 |
| | Proportion of polar atoms | 41.463 |
| | Alpha sphere density | 6.007 |
| | Cent. of mass - Alpha Sphere max dist | 16.279 |
| | Flexibility | 0.982 |
| Pocket 5 | | |
| | Score | 0.047 |
| | Druggability Score | 0.001 |
| | Number of Alpha Spheres | 16.000 |
| | Total SASA | 24.021 |
| | Polar SASA | 10.737 |
| | Apolar SASA | 13.284 |
| | Volume | 89.878 |
| | Mean local hydrophobic density | 6.000 |
| | Mean alpha sphere radius | 3.567 |
| | Mean alp. sph. solvent access | 0.402 |
| | Apolar alpha sphere proportion | 0.438 |
| | Hydrophobicity score | 22.143 |
| | Volume score | 4.571 |
| | Polarity score | 4.000 |
| | Charge score | 0.000 |
| | Proportion of polar atoms | 42.857 |
| | Alpha sphere density | 1.646 |
| | Cent. of mass - Alpha Sphere max dist | 3.684 |
| | Flexibility | 0.663 |
| Pocket 6 | | |
| | Score | 0.045 |
| | Druggability Score | 0.002 |
| | Number of Alpha Spheres | 26.000 |
| | Total SASA | 35.891 |
| | Polar SASA | 12.946 |
| | Apolar SASA | 22.945 |
| | Volume | 131.564 |
| | Mean local hydrophobic density | 6.000 |
| | Mean alpha sphere radius | 3.700 |
| | Mean alp. sph. solvent access | 0.421 |
| | Apolar alpha sphere proportion | 0.269 |
| | Hydrophobicity score | 23.700 |
| | Volume score | 4.400 |
| | Polarity score | 5.000 |
| | Charge score | 0.000 |
| | Proportion of polar atoms | 38.889 |

| | | |
|---|---|---|
| | Alpha sphere density | 1.915 |
| | Cent. of mass - Alpha Sphere max dist | 5.526 |
| | Flexibility | 0.970 |
| Pocket 7 | | |
| | Score | 0.023 |
| | Druggability Score | 0.001 |
| | Number of Alpha Spheres | 18.000 |
| | Total SASA | 40.583 |
| | Polar SASA | 26.091 |
| | Apolar SASA | 14.492 |
| | Volume | 117.018 |
| | Mean local hydrophobic density | 2.000 |
| | Mean alpha sphere radius | 3.545 |
| | Mean alp. sph. solvent access | 0.367 |
| | Apolar alpha sphere proportion | 0.167 |
| | Hydrophobicity score | 25.500 |
| | Volume score | 5.125 |
| | Polarity score | 5.000 |
| | Charge score | 3.000 |
| | Proportion of polar atoms | 52.941 |
| | Alpha sphere density | 2.061 |
| | Cent. of mass - Alpha Sphere max dist | 4.677 |
| | Flexibility | 0.910 |
| Pocket 8 | | |
| | Score | 0.010 |
| | Druggability Score | 0.023 |
| | Number of Alpha Spheres | 28.000 |
| | Total SASA | 57.172 |
| | Polar SASA | 17.320 |
| | Apolar SASA | 39.852 |
| | Volume | 182.406 |
| | Mean local hydrophobic density | 19.000 |
| | Mean alpha sphere radius | 3.663 |
| | Mean alp. sph. solvent access | 0.419 |
| | Apolar alpha sphere proportion | 0.714 |
| | Hydrophobicity score | 5.714 |
| | Volume score | 4.429 |
| | Polarity score | 6.000 |
| | Charge score | -1.000 |
| | Proportion of polar atoms | 38.889 |
| | Alpha sphere density | 2.417 |
| | Cent. of mass - Alpha Sphere max dist | 5.293 |

| | | |
|---|---|---|
| | Flexibility | 0.721 |
| Pocket 9 | | |
| | Score | -0.003 |
| | Druggability Score | 0.001 |
| | Number of Alpha Spheres | 39.000 |
| | Total SASA | 120.430 |
| | Polar SASA | 92.654 |
| | Apolar SASA | 27.775 |
| | Volume | 430.069 |
| | Mean local hydrophobic density | 3.667 |
| | Mean alpha sphere radius | 3.811 |
| | Mean alp. sph. solvent access | 0.493 |
| | Apolar alpha sphere proportion | 0.154 |
| | Hydrophobicity score | 17.429 |
| | Volume score | 4.357 |
| | Polarity score | 8.000 |
| | Charge score | 1.000 |
| | Proportion of polar atoms | 53.571 |
| | Alpha sphere density | 4.628 |
| | Cent. of mass - Alpha Sphere max dist | 11.598 |
| | Flexibility | 0.975 |
| Pocket 10 | | |
| | Score | -0.004 |
| | Druggability Score | 0.002 |
| | Number of Alpha Spheres | 32.000 |
| | Total SASA | 99.213 |
| | Polar SASA | 66.607 |
| | Apolar SASA | 32.606 |
| | Volume | 310.538 |
| | Mean local hydrophobic density | 1.000 |
| | Mean alpha sphere radius | 3.711 |
| | Mean alp. sph. solvent access | 0.395 |
| | Apolar alpha sphere proportion | 0.062 |
| | Hydrophobicity score | 16.429 |
| | Volume score | 4.071 |
| | Polarity score | 9.000 |
| | Charge score | 0.000 |
| | Proportion of polar atoms | 60.000 |
| | Alpha sphere density | 3.904 |
| | Cent. of mass - Alpha Sphere max dist | 10.958 |
| | Flexibility | 0.935 |

Pocket 11

| | |
|---|---|
| Score | -0.011 |
| Druggability Score | 0.003 |
| Number of Alpha Spheres | 35.000 |
| Total SASA | 64.448 |
| Polar SASA | 25.804 |
| Apolar SASA | 38.644 |
| Volume | 177.237 |
| Mean local hydrophobic density | 4.667 |
| Mean alpha sphere radius | 3.704 |
| Mean alp. sph. solvent access | 0.460 |
| Apolar alpha sphere proportion | 0.171 |
| Hydrophobicity score | 45.462 |
| Volume score | 4.231 |
| Polarity score | 5.000 |
| Charge score | 1.000 |
| Proportion of polar atoms | 47.619 |
| Alpha sphere density | 2.336 |
| Cent. of mass - Alpha Sphere max dist | 7.087 |
| Flexibility | 0.951 |

Pocket 12

| | |
|---|---|
| Score | -0.012 |
| Druggability Score | 0.004 |
| Number of Alpha Spheres | 39.000 |
| Total SASA | 127.083 |
| Polar SASA | 72.739 |
| Apolar SASA | 54.343 |
| Volume | 336.074 |
| Mean local hydrophobic density | 7.778 |
| Mean alpha sphere radius | 3.932 |
| Mean alp. sph. solvent access | 0.432 |
| Apolar alpha sphere proportion | 0.231 |
| Hydrophobicity score | 23.769 |
| Volume score | 4.385 |
| Polarity score | 8.000 |
| Charge score | 4.000 |
| Proportion of polar atoms | 46.154 |
| Alpha sphere density | 4.882 |
| Cent. of mass - Alpha Sphere max dist | 10.925 |
| Flexibility | 0.980 |

Pocket 13

| | |
|---|---|
| Score | -0.014 |

| | |
|---|---|
| Druggability Score | 0.009 |
| Number of Alpha Spheres | 25.000 |
| Total SASA | 47.080 |
| Polar SASA | 9.643 |
| Apolar SASA | 37.436 |
| Volume | 132.102 |
| Mean local hydrophobic density | 15.000 |
| Mean alpha sphere radius | 3.775 |
| Mean alp. sph. solvent access | 0.442 |
| Apolar alpha sphere proportion | 0.640 |
| Hydrophobicity score | 11.667 |
| Volume score | 4.444 |
| Polarity score | 6.000 |
| Charge score | 1.000 |
| Proportion of polar atoms | 41.176 |
| Alpha sphere density | 1.799 |
| Cent. of mass - Alpha Sphere max dist | 3.916 |
| Flexibility | 0.847 |

Pocket 14

| | |
|---|---|
| Score | -0.016 |
| Druggability Score | 0.000 |
| Number of Alpha Spheres | 22.000 |
| Total SASA | 44.894 |
| Polar SASA | 29.195 |
| Apolar SASA | 15.699 |
| Volume | 131.939 |
| Mean local hydrophobic density | 1.000 |
| Mean alpha sphere radius | 3.725 |
| Mean alp. sph. solvent access | 0.487 |
| Apolar alpha sphere proportion | 0.091 |
| Hydrophobicity score | 6.222 |
| Volume score | 3.778 |
| Polarity score | 7.000 |
| Charge score | -2.000 |
| Proportion of polar atoms | 47.059 |
| Alpha sphere density | 1.726 |
| Cent. of mass - Alpha Sphere max dist | 3.307 |
| Flexibility | 0.662 |

Pocket 15

| | |
|---|---|
| Score | -0.019 |
| Druggability Score | 0.005 |
| Number of Alpha Spheres | 33.000 |

| | |
|---|---|
| Total SASA | 95.612 |
| Polar SASA | 55.760 |
| Apolar SASA | 39.852 |
| Volume | 341.750 |
| Mean local hydrophobic density | 15.000 |
| Mean alpha sphere radius | 3.953 |
| Mean alp. sph. solvent access | 0.546 |
| Apolar alpha sphere proportion | 0.485 |
| Hydrophobicity score | 20.273 |
| Volume score | 3.909 |
| Polarity score | 6.000 |
| Charge score | 1.000 |
| Proportion of polar atoms | 37.500 |
| Alpha sphere density | 3.561 |
| Cent. of mass - Alpha Sphere max dist | 8.618 |
| Flexibility | 0.955 |

Pocket 16

| | |
|---|---|
| Score | -0.019 |
| Druggability Score | 0.001 |
| Number of Alpha Spheres | 61.000 |
| Total SASA | 128.326 |
| Polar SASA | 68.994 |
| Apolar SASA | 59.332 |
| Volume | 425.414 |
| Mean local hydrophobic density | 3.000 |
| Mean alpha sphere radius | 3.987 |
| Mean alp. sph. solvent access | 0.561 |
| Apolar alpha sphere proportion | 0.066 |
| Hydrophobicity score | 11.417 |
| Volume score | 3.500 |
| Polarity score | 7.000 |
| Charge score | -1.000 |
| Proportion of polar atoms | 45.946 |
| Alpha sphere density | 4.254 |
| Cent. of mass - Alpha Sphere max dist | 9.492 |
| Flexibility | 0.860 |

Pocket 17

| | |
|---|---|
| Score | -0.020 |
| Druggability Score | 0.000 |
| Number of Alpha Spheres | 16.000 |
| Total SASA | 23.590 |
| Polar SASA | 13.929 |

| | | |
|---|---|---|
| | Apolar SASA | 9.661 |
| | Volume | 67.593 |
| | Mean local hydrophobic density | 3.000 |
| | Mean alpha sphere radius | 3.601 |
| | Mean alp. sph. solvent access | 0.406 |
| | Apolar alpha sphere proportion | 0.250 |
| | Hydrophobicity score | 21.375 |
| | Volume score | 4.625 |
| | Polarity score | 6.000 |
| | Charge score | -1.000 |
| | Proportion of polar atoms | 50.000 |
| | Alpha sphere density | 0.899 |
| | Cent. of mass - Alpha Sphere max dist | 2.064 |
| | Flexibility | 0.952 |
| Pocket 18 | | |
| | Score | -0.023 |
| | Druggability Score | 0.003 |
| | Number of Alpha Spheres | 41.000 |
| | Total SASA | 160.190 |
| | Polar SASA | 93.770 |
| | Apolar SASA | 66.420 |
| | Volume | 544.982 |
| | Mean local hydrophobic density | 5.333 |
| | Mean alpha sphere radius | 3.970 |
| | Mean alp. sph. solvent access | 0.566 |
| | Apolar alpha sphere proportion | 0.220 |
| | Hydrophobicity score | 8.688 |
| | Volume score | 3.438 |
| | Polarity score | 9.000 |
| | Charge score | -1.000 |
| | Proportion of polar atoms | 52.941 |
| | Alpha sphere density | 6.182 |
| | Cent. of mass - Alpha Sphere max dist | 16.326 |
| | Flexibility | 0.683 |
| Pocket 19 | | |
| | Score | -0.025 |
| | Druggability Score | 0.017 |
| | Number of Alpha Spheres | 33.000 |
| | Total SASA | 127.982 |
| | Polar SASA | 49.486 |
| | Apolar SASA | 78.496 |
| | Volume | 489.355 |

| | | |
|---|---|---|
| | Mean local hydrophobic density | 13.000 |
| | Mean alpha sphere radius | 3.987 |
| | Mean alp. sph. solvent access | 0.524 |
| | Apolar alpha sphere proportion | 0.424 |
| | Hydrophobicity score | 20.778 |
| | Volume score | 2.778 |
| | Polarity score | 2.000 |
| | Charge score | 2.000 |
| | Proportion of polar atoms | 45.833 |
| | Alpha sphere density | 4.765 |
| | Cent. of mass - Alpha Sphere max dist | 10.461 |
| | Flexibility | 0.425 |
| Pocket 20 | | |
| | Score | -0.030 |
| | Druggability Score | 0.022 |
| | Number of Alpha Spheres | 37.000 |
| | Total SASA | 90.221 |
| | Polar SASA | 43.124 |
| | Apolar SASA | 47.097 |
| | Volume | 252.489 |
| | Mean local hydrophobic density | 11.000 |
| | Mean alpha sphere radius | 3.864 |
| | Mean alp. sph. solvent access | 0.429 |
| | Apolar alpha sphere proportion | 0.324 |
| | Hydrophobicity score | -1.231 |
| | Volume score | 3.154 |
| | Polarity score | 9.000 |
| | Charge score | -2.000 |
| | Proportion of polar atoms | 54.167 |
| | Alpha sphere density | 3.172 |
| | Cent. of mass - Alpha Sphere max dist | 8.628 |
| | Flexibility | 0.921 |
| Pocket 21 | | |
| | Score | -0.031 |
| | Druggability Score | 0.002 |
| | Number of Alpha Spheres | 22.000 |
| | Total SASA | 54.209 |
| | Polar SASA | 22.810 |
| | Apolar SASA | 31.398 |
| | Volume | 174.297 |
| | Mean local hydrophobic density | 7.000 |
| | Mean alpha sphere radius | 3.758 |

| | | |
|---|---|---|
| | Mean alp. sph. solvent access | 0.463 |
| | Apolar alpha sphere proportion | 0.364 |
| | Hydrophobicity score | 12.500 |
| | Volume score | 4.375 |
| | Polarity score | 5.000 |
| | Charge score | 3.000 |
| | Proportion of polar atoms | 47.059 |
| | Alpha sphere density | 1.982 |
| | Cent. of mass - Alpha Sphere max dist | 5.074 |
| | Flexibility | 0.959 |
| Pocket 22 | | |
| | Score | -0.031 |
| | Druggability Score | 0.011 |
| | Number of Alpha Spheres | 25.000 |
| | Total SASA | 130.159 |
| | Polar SASA | 55.286 |
| | Apolar SASA | 74.873 |
| | Volume | 428.973 |
| | Mean local hydrophobic density | 8.000 |
| | Mean alpha sphere radius | 4.123 |
| | Mean alp. sph. solvent access | 0.613 |
| | Apolar alpha sphere proportion | 0.400 |
| | Hydrophobicity score | 9.556 |
| | Volume score | 3.444 |
| | Polarity score | 4.000 |
| | Charge score | 3.000 |
| | Proportion of polar atoms | 41.667 |
| | Alpha sphere density | 5.111 |
| | Cent. of mass - Alpha Sphere max dist | 12.705 |
| | Flexibility | 0.831 |
| Pocket 23 | | |
| | Score | -0.035 |
| | Druggability Score | 0.000 |
| | Number of Alpha Spheres | 27.000 |
| | Total SASA | 121.655 |
| | Polar SASA | 72.142 |
| | Apolar SASA | 49.513 |
| | Volume | 507.733 |
| | Mean local hydrophobic density | 0.000 |
| | Mean alpha sphere radius | 4.125 |
| | Mean alp. sph. solvent access | 0.597 |
| | Apolar alpha sphere proportion | 0.037 |

| | |
|---|---|
| Hydrophobicity score | 10.111 |
| Volume score | 4.111 |
| Polarity score | 7.000 |
| Charge score | -2.000 |
| Proportion of polar atoms | 56.522 |
| Alpha sphere density | 4.643 |
| Cent. of mass - Alpha Sphere max dist | 12.729 |
| Flexibility | 0.904 |

Pocket 24

| | |
|---|---|
| Score | -0.041 |
| Druggability Score | 0.010 |
| Number of Alpha Spheres | 20.000 |
| Total SASA | 83.737 |
| Polar SASA | 29.394 |
| Apolar SASA | 54.343 |
| Volume | 323.576 |
| Mean local hydrophobic density | 13.733 |
| Mean alpha sphere radius | 4.132 |
| Mean alp. sph. solvent access | 0.691 |
| Apolar alpha sphere proportion | 0.750 |
| Hydrophobicity score | 54.000 |
| Volume score | 6.000 |
| Polarity score | 3.000 |
| Charge score | 1.000 |
| Proportion of polar atoms | 33.333 |
| Alpha sphere density | 3.203 |
| Cent. of mass - Alpha Sphere max dist | 8.473 |
| Flexibility | 0.579 |

Pocket 25

| | |
|---|---|
| Score | -0.042 |
| Druggability Score | 0.000 |
| Number of Alpha Spheres | 15.000 |
| Total SASA | 71.437 |
| Polar SASA | 55.738 |
| Apolar SASA | 15.699 |
| Volume | 173.548 |
| Mean local hydrophobic density | 0.000 |
| Mean alpha sphere radius | 3.690 |
| Mean alp. sph. solvent access | 0.641 |
| Apolar alpha sphere proportion | 0.000 |
| Hydrophobicity score | -4.000 |
| Volume score | 3.400 |

| | | |
|---|---|---|
| | Polarity score | 3.000 |
| | Charge score | 1.000 |
| | Proportion of polar atoms | 53.333 |
| | Alpha sphere density | 2.763 |
| | Cent. of mass - Alpha Sphere max dist | 5.657 |
| | Flexibility | 0.045 |
| Pocket 26 | | |
| | Score | -0.045 |
| | Druggability Score | 0.000 |
| | Number of Alpha Spheres | 15.000 |
| | Total SASA | 87.649 |
| | Polar SASA | 55.043 |
| | Apolar SASA | 32.606 |
| | Volume | 307.814 |
| | Mean local hydrophobic density | 0.000 |
| | Mean alpha sphere radius | 3.923 |
| | Mean alp. sph. solvent access | 0.480 |
| | Apolar alpha sphere proportion | 0.067 |
| | Hydrophobicity score | -16.500 |
| | Volume score | 4.100 |
| | Polarity score | 8.000 |
| | Charge score | 0.000 |
| | Proportion of polar atoms | 68.750 |
| | Alpha sphere density | 3.426 |
| | Cent. of mass - Alpha Sphere max dist | 7.776 |
| | Flexibility | 0.805 |
| Pocket 27 | | |
| | Score | -0.046 |
| | Druggability Score | 0.001 |
| | Number of Alpha Spheres | 15.000 |
| | Total SASA | 65.284 |
| | Polar SASA | 31.471 |
| | Apolar SASA | 33.814 |
| | Volume | 186.697 |
| | Mean local hydrophobic density | 6.000 |
| | Mean alpha sphere radius | 3.805 |
| | Mean alp. sph. solvent access | 0.609 |
| | Apolar alpha sphere proportion | 0.467 |
| | Hydrophobicity score | 13.333 |
| | Volume score | 5.000 |
| | Polarity score | 3.000 |
| | Charge score | 1.000 |

| | |
|---|---|
| Proportion of polar atoms | 35.294 |
| Alpha sphere density | 2.477 |
| Cent. of mass - Alpha Sphere max dist | 6.343 |
| Flexibility | 0.997 |

Pocket 28

| | |
|---|---|
| Score | -0.048 |
| Druggability Score | 0.417 |
| Number of Alpha Spheres | 69.000 |
| Total SASA | 155.534 |
| Polar SASA | 61.339 |
| Apolar SASA | 94.195 |
| Volume | 467.668 |
| Mean local hydrophobic density | 31.029 |
| Mean alpha sphere radius | 3.908 |
| Mean alp. sph. solvent access | 0.496 |
| Apolar alpha sphere proportion | 0.507 |
| Hydrophobicity score | 10.000 |
| Volume score | 3.867 |
| Polarity score | 8.000 |
| Charge score | 1.000 |
| Proportion of polar atoms | 34.146 |
| Alpha sphere density | 4.860 |
| Cent. of mass - Alpha Sphere max dist | 10.044 |
| Flexibility | 0.938 |

Pocket 29

| | |
|---|---|
| Score | -0.048 |
| Druggability Score | 0.018 |
| Number of Alpha Spheres | 53.000 |
| Total SASA | 127.865 |
| Polar SASA | 51.784 |
| Apolar SASA | 76.081 |
| Volume | 424.183 |
| Mean local hydrophobic density | 12.714 |
| Mean alpha sphere radius | 3.896 |
| Mean alp. sph. solvent access | 0.467 |
| Apolar alpha sphere proportion | 0.264 |
| Hydrophobicity score | -7.750 |
| Volume score | 3.917 |
| Polarity score | 9.000 |
| Charge score | 1.000 |
| Proportion of polar atoms | 43.750 |
| Alpha sphere density | 4.109 |

| | | |
|---|---|---|
| | Cent. of mass - Alpha Sphere max dist | 12.084 |
| | Flexibility | 0.800 |
| Pocket 30 | | |
| | Score | -0.053 |
| | Druggability Score | 0.000 |
| | Number of Alpha Spheres | 17.000 |
| | Total SASA | 48.653 |
| | Polar SASA | 28.123 |
| | Apolar SASA | 20.530 |
| | Volume | 185.780 |
| | Mean local hydrophobic density | 6.000 |
| | Mean alpha sphere radius | 3.997 |
| | Mean alp. sph. solvent access | 0.631 |
| | Apolar alpha sphere proportion | 0.412 |
| | Hydrophobicity score | 13.500 |
| | Volume score | 5.167 |
| | Polarity score | 5.000 |
| | Charge score | 1.000 |
| | Proportion of polar atoms | 46.154 |
| | Alpha sphere density | 1.611 |
| | Cent. of mass - Alpha Sphere max dist | 3.107 |
| | Flexibility | 0.887 |
| Pocket 31 | | |
| | Score | -0.065 |
| | Druggability Score | 0.000 |
| | Number of Alpha Spheres | 37.000 |
| | Total SASA | 118.953 |
| | Polar SASA | 68.232 |
| | Apolar SASA | 50.720 |
| | Volume | 407.559 |
| | Mean local hydrophobic density | 3.000 |
| | Mean alpha sphere radius | 3.961 |
| | Mean alp. sph. solvent access | 0.574 |
| | Apolar alpha sphere proportion | 0.108 |
| | Hydrophobicity score | -5.385 |
| | Volume score | 4.615 |
| | Polarity score | 12.000 |
| | Charge score | 4.000 |
| | Proportion of polar atoms | 62.069 |
| | Alpha sphere density | 3.982 |
| | Cent. of mass - Alpha Sphere max dist | 11.189 |
| | Flexibility | 0.868 |

Pocket 32

| | |
|---|---|
| Score | -0.069 |
| Druggability Score | 0.007 |
| Number of Alpha Spheres | 30.000 |
| Total SASA | 103.317 |
| Polar SASA | 35.690 |
| Apolar SASA | 67.627 |
| Volume | 311.939 |
| Mean local hydrophobic density | 10.667 |
| Mean alpha sphere radius | 3.863 |
| Mean alp. sph. solvent access | 0.514 |
| Apolar alpha sphere proportion | 0.400 |
| Hydrophobicity score | 13.455 |
| Volume score | 3.545 |
| Polarity score | 3.000 |
| Charge score | 0.000 |
| Proportion of polar atoms | 37.500 |
| Alpha sphere density | 3.418 |
| Cent. of mass - Alpha Sphere max dist | 8.972 |
| Flexibility | 0.532 |

Pocket 33

| | |
|---|---|
| Score | -0.069 |
| Druggability Score | 0.001 |
| Number of Alpha Spheres | 19.000 |
| Total SASA | 69.871 |
| Polar SASA | 31.227 |
| Apolar SASA | 38.644 |
| Volume | 228.409 |
| Mean local hydrophobic density | 6.000 |
| Mean alpha sphere radius | 3.936 |
| Mean alp. sph. solvent access | 0.537 |
| Apolar alpha sphere proportion | 0.368 |
| Hydrophobicity score | 23.167 |
| Volume score | 3.500 |
| Polarity score | 3.000 |
| Charge score | 0.000 |
| Proportion of polar atoms | 43.750 |
| Alpha sphere density | 2.306 |
| Cent. of mass - Alpha Sphere max dist | 5.844 |
| Flexibility | 0.697 |

Pocket 34

| | |
|---|---|
| Score | -0.083 |
| Druggability Score | 0.002 |
| Number of Alpha Spheres | 19.000 |
| Total SASA | 62.900 |
| Polar SASA | 22.810 |
| Apolar SASA | 40.089 |
| Volume | 216.789 |
| Mean local hydrophobic density | 11.000 |
| Mean alpha sphere radius | 3.956 |
| Mean alp. sph. solvent access | 0.533 |
| Apolar alpha sphere proportion | 0.632 |
| Hydrophobicity score | 38.333 |
| Volume score | 4.000 |
| Polarity score | 2.000 |
| Charge score | 1.000 |
| Proportion of polar atoms | 35.714 |
| Alpha sphere density | 1.881 |
| Cent. of mass - Alpha Sphere max dist | 5.156 |
| Flexibility | 0.584 |

Pocket 35

| | |
|---|---|
| Score | -0.084 |
| Druggability Score | 0.005 |
| Number of Alpha Spheres | 58.000 |
| Total SASA | 160.254 |
| Polar SASA | 116.780 |
| Apolar SASA | 43.475 |
| Volume | 456.786 |
| Mean local hydrophobic density | 5.750 |
| Mean alpha sphere radius | 3.785 |
| Mean alp. sph. solvent access | 0.475 |
| Apolar alpha sphere proportion | 0.138 |
| Hydrophobicity score | 6.059 |
| Volume score | 4.059 |
| Polarity score | 11.000 |
| Charge score | 2.000 |
| Proportion of polar atoms | 56.098 |
| Alpha sphere density | 4.854 |
| Cent. of mass - Alpha Sphere max dist | 14.097 |
| Flexibility | 0.833 |

Pocket 36

| | |
|---|---|
| Score | -0.094 |
| Druggability Score | 0.000 |

| | |
|---|---|
| Number of Alpha Spheres | 15.000 |
| Total SASA | 76.028 |
| Polar SASA | 39.799 |
| Apolar SASA | 36.229 |
| Volume | 219.278 |
| Mean local hydrophobic density | 0.000 |
| Mean alpha sphere radius | 4.016 |
| Mean alp. sph. solvent access | 0.546 |
| Apolar alpha sphere proportion | 0.067 |
| Hydrophobicity score | 11.714 |
| Volume score | 3.429 |
| Polarity score | 4.000 |
| Charge score | 0.000 |
| Proportion of polar atoms | 57.143 |
| Alpha sphere density | 2.401 |
| Cent. of mass - Alpha Sphere max dist | 5.719 |
| Flexibility | 0.544 |

Pocket 37

| | |
|---|---|
| Score | -0.098 |
| Druggability Score | 0.000 |
| Number of Alpha Spheres | 16.000 |
| Total SASA | 87.516 |
| Polar SASA | 54.910 |
| Apolar SASA | 32.606 |
| Volume | 254.181 |
| Mean local hydrophobic density | 1.000 |
| Mean alpha sphere radius | 3.871 |
| Mean alp. sph. solvent access | 0.588 |
| Apolar alpha sphere proportion | 0.125 |
| Hydrophobicity score | -4.714 |
| Volume score | 4.000 |
| Polarity score | 6.000 |
| Charge score | 1.000 |
| Proportion of polar atoms | 53.333 |
| Alpha sphere density | 2.837 |
| Cent. of mass - Alpha Sphere max dist | 7.218 |
| Flexibility | 0.814 |

Pocket 38

| | |
|---|---|
| Score | -0.098 |
| Druggability Score | 0.222 |
| Number of Alpha Spheres | 60.000 |
| Total SASA | 162.716 |

| | |
|---|---|
| Polar SASA | 81.488 |
| Apolar SASA | 81.228 |
| Volume | 633.485 |
| Mean local hydrophobic density | 29.355 |
| Mean alpha sphere radius | 3.915 |
| Mean alp. sph. solvent access | 0.521 |
| Apolar alpha sphere proportion | 0.517 |
| Hydrophobicity score | 22.812 |
| Volume score | 4.625 |
| Polarity score | 9.000 |
| Charge score | 3.000 |
| Proportion of polar atoms | 37.209 |
| Alpha sphere density | 4.558 |
| Cent. of mass - Alpha Sphere max dist | 10.708 |
| Flexibility | 0.733 |

Pocket 39

| | |
|---|---|
| Score | -0.099 |
| Druggability Score | 0.000 |
| Number of Alpha Spheres | 23.000 |
| Total SASA | 78.389 |
| Polar SASA | 54.237 |
| Apolar SASA | 24.153 |
| Volume | 209.590 |
| Mean local hydrophobic density | 1.000 |
| Mean alpha sphere radius | 3.890 |
| Mean alp. sph. solvent access | 0.492 |
| Apolar alpha sphere proportion | 0.087 |
| Hydrophobicity score | 26.714 |
| Volume score | 4.286 |
| Polarity score | 5.000 |
| Charge score | 1.000 |
| Proportion of polar atoms | 58.824 |
| Alpha sphere density | 2.264 |
| Cent. of mass - Alpha Sphere max dist | 5.355 |
| Flexibility | 0.970 |

Pocket 40

| | |
|---|---|
| Score | -0.101 |
| Druggability Score | 0.000 |
| Number of Alpha Spheres | 16.000 |
| Total SASA | 79.534 |
| Polar SASA | 52.966 |
| Apolar SASA | 26.568 |

| | | |
|---|---|---|
| | Volume | 190.175 |
| | Mean local hydrophobic density | 2.000 |
| | Mean alpha sphere radius | 3.898 |
| | Mean alp. sph. solvent access | 0.512 |
| | Apolar alpha sphere proportion | 0.188 |
| | Hydrophobicity score | 15.833 |
| | Volume score | 5.000 |
| | Polarity score | 5.000 |
| | Charge score | 2.000 |
| | Proportion of polar atoms | 60.000 |
| | Alpha sphere density | 2.471 |
| | Cent. of mass - Alpha Sphere max dist | 5.287 |
| | Flexibility | 0.462 |
| Pocket 41 | | |
| | Score | -0.106 |
| | Druggability Score | 0.014 |
| | Number of Alpha Spheres | 25.000 |
| | Total SASA | 87.337 |
| | Polar SASA | 18.502 |
| | Apolar SASA | 68.835 |
| | Volume | 282.314 |
| | Mean local hydrophobic density | 18.000 |
| | Mean alpha sphere radius | 4.009 |
| | Mean alp. sph. solvent access | 0.608 |
| | Apolar alpha sphere proportion | 0.760 |
| | Hydrophobicity score | 36.200 |
| | Volume score | 5.000 |
| | Polarity score | 3.000 |
| | Charge score | 0.000 |
| | Proportion of polar atoms | 23.810 |
| | Alpha sphere density | 2.544 |
| | Cent. of mass - Alpha Sphere max dist | 6.396 |
| | Flexibility | 0.903 |
| Pocket 42 | | |
| | Score | -0.112 |
| | Druggability Score | 0.017 |
| | Number of Alpha Spheres | 43.000 |
| | Total SASA | 121.716 |
| | Polar SASA | 51.673 |
| | Apolar SASA | 70.042 |
| | Volume | 411.515 |
| | Mean local hydrophobic density | 14.000 |

| | |
|---|---|
| Mean alpha sphere radius | 4.059 |
| Mean alp. sph. solvent access | 0.509 |
| Apolar alpha sphere proportion | 0.349 |
| Hydrophobicity score | 20.500 |
| Volume score | 4.000 |
| Polarity score | 8.000 |
| Charge score | 1.000 |
| Proportion of polar atoms | 44.828 |
| Alpha sphere density | 3.466 |
| Cent. of mass - Alpha Sphere max dist | 9.818 |
| Flexibility | 0.942 |

Pocket 43

| | |
|---|---|
| Score | -0.113 |
| Druggability Score | 0.001 |
| Number of Alpha Spheres | 44.000 |
| Total SASA | 136.831 |
| Polar SASA | 76.450 |
| Apolar SASA | 60.381 |
| Volume | 464.167 |
| Mean local hydrophobic density | 9.000 |
| Mean alpha sphere radius | 3.956 |
| Mean alp. sph. solvent access | 0.535 |
| Apolar alpha sphere proportion | 0.227 |
| Hydrophobicity score | -4.200 |
| Volume score | 4.500 |
| Polarity score | 8.000 |
| Charge score | 0.000 |
| Proportion of polar atoms | 48.485 |
| Alpha sphere density | 4.064 |
| Cent. of mass - Alpha Sphere max dist | 12.052 |
| Flexibility | 0.928 |

Pocket 44

| | |
|---|---|
| Score | -0.118 |
| Druggability Score | 0.187 |
| Number of Alpha Spheres | 41.000 |
| Total SASA | 155.589 |
| Polar SASA | 54.148 |
| Apolar SASA | 101.441 |
| Volume | 530.394 |
| Mean local hydrophobic density | 22.160 |
| Mean alpha sphere radius | 3.940 |
| Mean alp. sph. solvent access | 0.509 |

| | | |
|---|---|---|
| | Apolar alpha sphere proportion | 0.610 |
| | Hydrophobicity score | 7.100 |
| | Volume score | 4.300 |
| | Polarity score | 8.000 |
| | Charge score | 2.000 |
| | Proportion of polar atoms | 35.484 |
| | Alpha sphere density | 4.749 |
| | Cent. of mass - Alpha Sphere max dist | 12.554 |
| | Flexibility | 0.967 |
| Pocket 45 | | |
| | Score | -0.121 |
| | Druggability Score | 0.002 |
| | Number of Alpha Spheres | 15.000 |
| | Total SASA | 115.627 |
| | Polar SASA | 54.038 |
| | Apolar SASA | 61.589 |
| | Volume | 349.232 |
| | Mean local hydrophobic density | 8.000 |
| | Mean alpha sphere radius | 4.095 |
| | Mean alp. sph. solvent access | 0.698 |
| | Apolar alpha sphere proportion | 0.600 |
| | Hydrophobicity score | 5.750 |
| | Volume score | 4.500 |
| | Polarity score | 5.000 |
| | Charge score | 1.000 |
| | Proportion of polar atoms | 43.478 |
| | Alpha sphere density | 3.838 |
| | Cent. of mass - Alpha Sphere max dist | 8.174 |
| | Flexibility | 0.959 |
| Pocket 46 | | |
| | Score | -0.121 |
| | Druggability Score | 0.000 |
| | Number of Alpha Spheres | 21.000 |
| | Total SASA | 92.135 |
| | Polar SASA | 36.585 |
| | Apolar SASA | 55.551 |
| | Volume | 259.709 |
| | Mean local hydrophobic density | 1.000 |
| | Mean alpha sphere radius | 3.924 |
| | Mean alp. sph. solvent access | 0.573 |
| | Apolar alpha sphere proportion | 0.095 |
| | Hydrophobicity score | 3.625 |

| | | |
|---|---|---|
| | Volume score | 3.125 |
| | Polarity score | 4.000 |
| | Charge score | -2.000 |
| | Proportion of polar atoms | 47.368 |
| | Alpha sphere density | 2.617 |
| | Cent. of mass - Alpha Sphere max dist | 5.316 |
| | Flexibility | 0.577 |
| Pocket 47 | | |
| | Score | -0.129 |
| | Druggability Score | 0.003 |
| | Number of Alpha Spheres | 20.000 |
| | Total SASA | 81.657 |
| | Polar SASA | 32.144 |
| | Apolar SASA | 49.513 |
| | Volume | 192.281 |
| | Mean local hydrophobic density | 12.000 |
| | Mean alpha sphere radius | 3.802 |
| | Mean alp. sph. solvent access | 0.507 |
| | Apolar alpha sphere proportion | 0.650 |
| | Hydrophobicity score | 20.400 |
| | Volume score | 3.200 |
| | Polarity score | 1.000 |
| | Charge score | -1.000 |
| | Proportion of polar atoms | 33.333 |
| | Alpha sphere density | 2.173 |
| | Cent. of mass - Alpha Sphere max dist | 4.736 |
| | Flexibility | 0.882 |
| Pocket 48 | | |
| | Score | -0.132 |
| | Druggability Score | 0.008 |
| | Number of Alpha Spheres | 28.000 |
| | Total SASA | 117.142 |
| | Polar SASA | 59.196 |
| | Apolar SASA | 57.946 |
| | Volume | 305.573 |
| | Mean local hydrophobic density | 11.000 |
| | Mean alpha sphere radius | 3.982 |
| | Mean alp. sph. solvent access | 0.650 |
| | Apolar alpha sphere proportion | 0.429 |
| | Hydrophobicity score | 19.200 |
| | Volume score | 3.800 |
| | Polarity score | 5.000 |

| | | |
|---|---|---|
| | Charge score | 2.000 |
| | Proportion of polar atoms | 44.000 |
| | Alpha sphere density | 3.415 |
| | Cent. of mass - Alpha Sphere max dist | 10.477 |
| | Flexibility | 0.926 |
| Pocket 49 | | |
| | Score | -0.132 |
| | Druggability Score | 0.006 |
| | Number of Alpha Spheres | 23.000 |
| | Total SASA | 81.444 |
| | Polar SASA | 36.762 |
| | Apolar SASA | 44.682 |
| | Volume | 185.406 |
| | Mean local hydrophobic density | 15.000 |
| | Mean alpha sphere radius | 3.930 |
| | Mean alp. sph. solvent access | 0.476 |
| | Apolar alpha sphere proportion | 0.696 |
| | Hydrophobicity score | 38.333 |
| | Volume score | 4.444 |
| | Polarity score | 3.000 |
| | Charge score | 1.000 |
| | Proportion of polar atoms | 35.294 |
| | Alpha sphere density | 2.043 |
| | Cent. of mass - Alpha Sphere max dist | 5.135 |
| | Flexibility | 0.930 |
| Pocket 50 | | |
| | Score | -0.140 |
| | Druggability Score | 0.013 |
| | Number of Alpha Spheres | 49.000 |
| | Total SASA | 193.174 |
| | Polar SASA | 82.073 |
| | Apolar SASA | 111.102 |
| | Volume | 596.227 |
| | Mean local hydrophobic density | 17.895 |
| | Mean alpha sphere radius | 3.911 |
| | Mean alp. sph. solvent access | 0.577 |
| | Apolar alpha sphere proportion | 0.388 |
| | Hydrophobicity score | 18.867 |
| | Volume score | 3.867 |
| | Polarity score | 8.000 |
| | Charge score | -1.000 |
| | Proportion of polar atoms | 44.737 |

| | | |
|---|---|---|
| | Alpha sphere density | 5.646 |
| | Cent. of mass - Alpha Sphere max dist | 12.261 |
| | Flexibility | 0.862 |
| Pocket 51 | | |
| | Score | -0.143 |
| | Druggability Score | 0.000 |
| | Number of Alpha Spheres | 22.000 |
| | Total SASA | 108.236 |
| | Polar SASA | 35.779 |
| | Apolar SASA | 72.458 |
| | Volume | 342.493 |
| | Mean local hydrophobic density | 4.000 |
| | Mean alpha sphere radius | 4.103 |
| | Mean alp. sph. solvent access | 0.511 |
| | Apolar alpha sphere proportion | 0.227 |
| | Hydrophobicity score | 12.571 |
| | Volume score | 4.571 |
| | Polarity score | 3.000 |
| | Charge score | 1.000 |
| | Proportion of polar atoms | 41.176 |
| | Alpha sphere density | 3.105 |
| | Cent. of mass - Alpha Sphere max dist | 6.645 |
| | Flexibility | 0.799 |
| Pocket 52 | | |
| | Score | -0.146 |
| | Druggability Score | 0.000 |
| | Number of Alpha Spheres | 39.000 |
| | Total SASA | 139.074 |
| | Polar SASA | 71.447 |
| | Apolar SASA | 67.627 |
| | Volume | 411.859 |
| | Mean local hydrophobic density | 5.000 |
| | Mean alpha sphere radius | 3.842 |
| | Mean alp. sph. solvent access | 0.438 |
| | Apolar alpha sphere proportion | 0.154 |
| | Hydrophobicity score | 12.929 |
| | Volume score | 3.714 |
| | Polarity score | 8.000 |
| | Charge score | 0.000 |
| | Proportion of polar atoms | 51.852 |
| | Alpha sphere density | 3.832 |
| | Cent. of mass - Alpha Sphere max dist | 7.323 |

| | |
|---|---|
| Flexibility | 0.923 |

Pocket 53

| | |
|---|---|
| Score | -0.147 |
| Druggability Score | 0.000 |
| Number of Alpha Spheres | 32.000 |
| Total SASA | 125.518 |
| Polar SASA | 84.459 |
| Apolar SASA | 41.059 |
| Volume | 345.878 |
| Mean local hydrophobic density | 2.000 |
| Mean alpha sphere radius | 3.802 |
| Mean alp. sph. solvent access | 0.469 |
| Apolar alpha sphere proportion | 0.094 |
| Hydrophobicity score | 1.182 |
| Volume score | 4.364 |
| Polarity score | 8.000 |
| Charge score | 3.000 |
| Proportion of polar atoms | 54.167 |
| Alpha sphere density | 3.489 |
| Cent. of mass - Alpha Sphere max dist | 7.513 |
| Flexibility | 0.814 |

Pocket 54

| | |
|---|---|
| Score | -0.153 |
| Druggability Score | 0.000 |
| Number of Alpha Spheres | 20.000 |
| Total SASA | 96.633 |
| Polar SASA | 59.771 |
| Apolar SASA | 36.862 |
| Volume | 238.236 |
| Mean local hydrophobic density | 1.000 |
| Mean alpha sphere radius | 3.906 |
| Mean alp. sph. solvent access | 0.597 |
| Apolar alpha sphere proportion | 0.100 |
| Hydrophobicity score | 3.833 |
| Volume score | 3.833 |
| Polarity score | 5.000 |
| Charge score | 1.000 |
| Proportion of polar atoms | 58.824 |
| Alpha sphere density | 2.530 |
| Cent. of mass - Alpha Sphere max dist | 4.823 |
| Flexibility | 0.718 |

Pocket 55

| | |
|---|---|
| Score | -0.154 |
| Druggability Score | 0.006 |
| Number of Alpha Spheres | 40.000 |
| Total SASA | 138.765 |
| Polar SASA | 82.006 |
| Apolar SASA | 56.759 |
| Volume | 505.517 |
| Mean local hydrophobic density | 15.000 |
| Mean alpha sphere radius | 4.094 |
| Mean alp. sph. solvent access | 0.447 |
| Apolar alpha sphere proportion | 0.400 |
| Hydrophobicity score | 4.556 |
| Volume score | 4.111 |
| Polarity score | 7.000 |
| Charge score | -1.000 |
| Proportion of polar atoms | 43.333 |
| Alpha sphere density | 3.685 |
| Cent. of mass - Alpha Sphere max dist | 8.359 |
| Flexibility | 0.968 |

Pocket 56

| | |
|---|---|
| Score | -0.159 |
| Druggability Score | 0.007 |
| Number of Alpha Spheres | 40.000 |
| Total SASA | 167.935 |
| Polar SASA | 64.079 |
| Apolar SASA | 103.856 |
| Volume | 496.651 |
| Mean local hydrophobic density | 15.143 |
| Mean alpha sphere radius | 3.871 |
| Mean alp. sph. solvent access | 0.546 |
| Apolar alpha sphere proportion | 0.525 |
| Hydrophobicity score | 26.900 |
| Volume score | 5.200 |
| Polarity score | 7.000 |
| Charge score | 2.000 |
| Proportion of polar atoms | 32.258 |
| Alpha sphere density | 4.927 |
| Cent. of mass - Alpha Sphere max dist | 9.689 |
| Flexibility | 0.860 |

Pocket 57

| | |
|---|---|
| Score | -0.173 |

| | |
|---|---|
| Druggability Score | 0.003 |
| Number of Alpha Spheres | 38.000 |
| Total SASA | 126.095 |
| Polar SASA | 76.583 |
| Apolar SASA | 49.513 |
| Volume | 348.897 |
| Mean local hydrophobic density | 12.000 |
| Mean alpha sphere radius | 4.140 |
| Mean alp. sph. solvent access | 0.409 |
| Apolar alpha sphere proportion | 0.342 |
| Hydrophobicity score | 10.556 |
| Volume score | 4.222 |
| Polarity score | 8.000 |
| Charge score | 0.000 |
| Proportion of polar atoms | 52.381 |
| Alpha sphere density | 3.188 |
| Cent. of mass - Alpha Sphere max dist | 9.370 |
| Flexibility | 0.843 |

Pocket 58

| | |
|---|---|
| Score | -0.179 |
| Druggability Score | 0.009 |
| Number of Alpha Spheres | 75.000 |
| Total SASA | 193.565 |
| Polar SASA | 107.269 |
| Apolar SASA | 86.296 |
| Volume | 630.895 |
| Mean local hydrophobic density | 18.000 |
| Mean alpha sphere radius | 4.004 |
| Mean alp. sph. solvent access | 0.491 |
| Apolar alpha sphere proportion | 0.253 |
| Hydrophobicity score | -0.571 |
| Volume score | 4.143 |
| Polarity score | 10.000 |
| Charge score | 4.000 |
| Proportion of polar atoms | 53.191 |
| Alpha sphere density | 4.782 |
| Cent. of mass - Alpha Sphere max dist | 12.088 |
| Flexibility | 0.895 |

Pocket 59

| | |
|---|---|
| Score | -0.194 |
| Druggability Score | 0.003 |
| Number of Alpha Spheres | 24.000 |

| | |
|---|---|
| Total SASA | 125.629 |
| Polar SASA | 39.887 |
| Apolar SASA | 85.742 |
| Volume | 347.772 |
| Mean local hydrophobic density | 10.000 |
| Mean alpha sphere radius | 4.064 |
| Mean alp. sph. solvent access | 0.586 |
| Apolar alpha sphere proportion | 0.458 |
| Hydrophobicity score | 21.000 |
| Volume score | 4.250 |
| Polarity score | 5.000 |
| Charge score | 1.000 |
| Proportion of polar atoms | 45.000 |
| Alpha sphere density | 3.278 |
| Cent. of mass - Alpha Sphere max dist | 8.997 |
| Flexibility | 0.899 |

Pocket 60

| | |
|---|---|
| Score | -0.209 |
| Druggability Score | 0.666t |
| Number of Alpha Spheres | 157.000 |
| Total SASA | 436.409 |
| Polar SASA | 147.786 |
| Apolar SASA | 288.623 |
| Volume | 1667.223 |
| Mean local hydrophobic density | 20.340 |
| Mean alpha sphere radius | 4.062 |
| Mean alp. sph. solvent access | 0.507 |
| Apolar alpha sphere proportion | 0.299 |
| Hydrophobicity score | 27.125 |
| Volume score | 4.229 |
| Polarity score | 29.000 |
| Charge score | 0.000 |
| Proportion of polar atoms | 39.796 |
| Alpha sphere density | 9.556 |
| Cent. of mass - Alpha Sphere max dist | 25.332 |
| Flexibility | 0.844 |

Pocket 61

| | |
|---|---|
| Score | -0.221 |
| Druggability Score | 0.001 |
| Number of Alpha Spheres | 19.000 |
| Total SASA | 98.818 |
| Polar SASA | 25.153 |

| | |
|---|---|
| Apolar SASA | 73.665 |
| Volume | 195.952 |
| Mean local hydrophobic density | 10.000 |
| Mean alpha sphere radius | 3.903 |
| Mean alp. sph. solvent access | 0.542 |
| Apolar alpha sphere proportion | 0.579 |
| Hydrophobicity score | 43.250 |
| Volume score | 5.500 |
| Polarity score | 2.000 |
| Charge score | 1.000 |
| Proportion of polar atoms | 33.333 |
| Alpha sphere density | 1.951 |
| Cent. of mass - Alpha Sphere max dist | 5.180 |
| Flexibility | 0.636 |

Pocket 62

| | |
|---|---|
| Score | -0.233 |
| Druggability Score | 0.004 |
| Number of Alpha Spheres | 46.000 |
| Total SASA | 181.659 |
| Polar SASA | 91.087 |
| Apolar SASA | 90.572 |
| Volume | 598.479 |
| Mean local hydrophobic density | 14.000 |
| Mean alpha sphere radius | 4.154 |
| Mean alp. sph. solvent access | 0.523 |
| Apolar alpha sphere proportion | 0.326 |
| Hydrophobicity score | 8.643 |
| Volume score | 4.643 |
| Polarity score | 10.000 |
| Charge score | 5.000 |
| Proportion of polar atoms | 52.941 |
| Alpha sphere density | 4.318 |
| Cent. of mass - Alpha Sphere max dist | 12.678 |
| Flexibility | 0.414 |

Pocket 63

| | |
|---|---|
| Score | -0.297 |
| Druggability Score | 0.001 |
| Number of Alpha Spheres | 233.000 |
| Total SASA | 643.190 |
| Polar SASA | 320.436 |
| Apolar SASA | 322.753 |
| Volume | 2208.325 |

| | | |
|---|---|---|
| | Mean local hydrophobic density | 27.644 |
| | Mean alpha sphere radius | 3.985 |
| | Mean alp. sph. solvent access | 0.459 |
| | Apolar alpha sphere proportion | 0.386 |
| | Hydrophobicity score | 3.833 |
| | Volume score | 4.119 |
| | Polarity score | 25.000 |
| | Charge score | 2.000 |
| | Proportion of polar atoms | 41.406 |
| | Alpha sphere density | 10.983 |
| | Cent. of mass - Alpha Sphere max dist | 26.566 |
| | Flexibility | 0.588 |
| Pocket 64 | | |
| | Score | -0.328 |
| | Druggability Score | 0.018 |
| | Number of Alpha Spheres | 197.000 |
| | Total SASA | 495.749 |
| | Polar SASA | 230.070 |
| | Apolar SASA | 265.678 |
| | Volume | 1990.677 |
| | Mean local hydrophobic density | 28.841 |
| | Mean alpha sphere radius | 3.927 |
| | Mean alp. sph. solvent access | 0.465 |
| | Apolar alpha sphere proportion | 0.350 |
| | Hydrophobicity score | 12.600 |
| | Volume score | 4.800 |
| | Polarity score | 23.000 |
| | Charge score | 5.000 |
| | Proportion of polar atoms | 39.806 |
| | Alpha sphere density | 8.845 |
| | Cent. of mass - Alpha Sphere max dist | 20.290 |
| | Flexibility | 0.879 |
